# Exact calculation of stationary solution and parameter sensitivity analysis of stochastic continuous time Boolean models

**DOI:** 10.1101/794230

**Authors:** Mihály Koltai, Vincent Noel, Andrei Zinovyev, Laurence Calzone, Emmanuel Barillot

## Abstract

**Motivation:** Solutions to stochastic Boolean models are usually estimated by Monte Carlo simulations, but as the state space of these models can be enormous, there is an inherent uncertainty about the accuracy of Monte Carlo estimates and whether simulations have reached all asymptotic solutions. Moreover, these models have timescale parameters (transition rates) that the probability values of stationary solutions depend on in complex ways that have not been analyzed yet in the literature. These two fundamental uncertainties call for an exact calculation method for this class of models.

**Results:** We show that the stationary probability values of the attractors of stochastic (asynchronous) continuous time Boolean models can be exactly calculated. The calculation does not require Monte Carlo simulations, instead it uses an exact matrix calculation method previously applied in the context of chemical kinetics. Using this approach, we also analyze the under-explored question of the effect of transition rates on the stationary solutions and show the latter can be sensitive to parameter changes. The analysis distinguishes processes that are robust or, alternatively, sensitive to parameter values, providing both methodological and biological insights.

**Supplementary information:** Supplementary data available at *bioRxiv* online.

**Availability and implementation:** The calculation method described in the article is available as the ExaStoLog MATLAB package on GitHub at https://github.com/sysbio-curie/exact-stoch-log-mod

## 1 Introduction

One of the principle aims of systems biology is to describe and understand the complex molecular networks that regulate the functioning of a cell (Alon, 2006) by quantitative models. To do so numerous mathematical and computational formalisms have been used in the past decades (Le Novere, 2015). These range from quantitative and mechanistic models (stochastic or deterministic chemical kinetics, spatial models such as reaction-diffusion models) that require the knowledge of numerous biophysical constants (Calzone *et al.*, 2018) to higher level, more qualitative models such as fuzzy logic (Morris *et al.*, 2011; Aldridge *et al.*, 2009) and Boolean models (Wynn *et al.*, 2012; Morris *et al.*, 2010) that describe functional dependencies, but not the details of the biophysical mechanisms. Boolean models have the advantage that interactions between a model’s variables (typically genes and/or proteins) only need to be qualitatively defined and that calculations for this class of models are generally much faster than with continuous models (Naldi *et al.*, 2018), such as ordinary differential equations (ODE). Traditionally, Boolean modeling has been mainly used as a more qualitative approach, for quickly identifying the stationary states (attractors) of a particular model and test their robustness to initial conditions and/or perturbations. In most Boolean modeling platforms (Naldi *et al.*, 2018; Müssel *et al.*, 2010; Terfve *et al.*, 2012), time is described in discrete steps and the output of calculations are binary.

In recent years, efforts were made to bridge the gap between qualitative and quantitative modeling by a continuous time stochastic version of Boolean modeling (Stoll *et al.*, 2012, 2017). With this approach, the temporal evolution of a system is described as a continuous time Markov process on a Boolean state space. Mathematically, this is equivalent to a master equation that is a set of ODEs on the probability of the model’s space. In the MaBoSS modeling tool and its applications (Montagud *et al.*, 2017; Letort *et al.*, 2018; Béal *et al.*, 2019), this master equation is simulated by a kinetic Monte Carlo (Gillespie) algorithm (Rao and Arkin, 2003), as in chemical kinetics.

With this hybrid approach, we obtain continuous values for the transient and stationary probabilities of the states of a Boolean model, while still specifying only logical rules instead of chemical reactions with kinetic parameters. However, for the model’s continuous treatment, it is necessary to define transition rates for each of the model’s variables (nodes) that determine the probability of that variable being updated from 0 to 1 or *vice versa* at a given updating step of the Gillespie algorithm. Using the Gillespie algorithm enables the simulation of large logical models (Béal *et al.*, 2019), up to 200 or even more nodes.

With an increasing model size, the number of sample trajectories from which the probabilities are calculated is typically decreased in the interest of computation speed. Besides compromising the accuracy of probability estimates, this raises the issue of attractor reachability. As the state space of a Boolean model of *n* variables has a dimension of 2^*n*^, a limited number of stochastic simulations might not reach all attractors, leaving an uncertainty if the model’s behavior is fully explored with a given number of sample trajectories and simulation length.

Another question that needs to be addressed is parameter dependence, i.e., the dependence of the probability values of attractors on the transition rates. The probability value of an attractor state depends in complex ways on the transition rates along its path within the model’s state transition graph (STG), which is a directed graph of all model states with edges representing the possible transitions among them, as defined by the logical rules.

In the studies using the continuous time stochastic Boolean formalism, transition rates are usually assigned a default value (typically 1) for the sake of simplicity and minimizing parameter-dependent bias. However, transition rates are not neutral parameters: as shown in (Béal *et al.*, 2019), setting the value of transition rates based on expression data can improve a model’s predictive performance. This suggests that systematic parameter sensitivity analysis, similar to what is done for ODE models (Zi, 2011; Fröhlich *et al.*, 2017), can provide valuable insights on what the key variables and corresponding transition rates of a particular model are with regard to the probability of its output (phenotype) variables or its attractor states. For instance, we might ask the question in the case of a model with the outputs of proliferation versus cell death if there are transition rates that dominate the decision point between these two outcomes.

We provide here an exact method adopting mathematical techniques previously applied in the context of deterministic chemical kinetics (ODEs) (Gunawardena, 2012; Karp *et al.*, 2012; Mirzaev and Gunawardena, 2013) to calculate the stationary solutions of stochastic continuous time Boolean models. We make use of the fact that the kinetic matrix of logical models is typically very sparse, so that the calculation is as fast as or faster than stochastic simulations up to an intermediate size of around 20 nodes, while being exact.

Because of the exact nature of the calculation, it is guaranteed to find all states of the stationary solutions and their probability values after convergence, eliminating the issue of choosing a sufficient amount of time and number of sample trajectories to reach convergence and all attractors of a model. We perform parameter sensitivity analysis and visualization of solutions and their dependence on parameters on a number of published Boolean models (Traynard *et al.*, 2016; Zañudo *et al.*, 2017; Sahin *et al.*, 2009; Cohen *et al.*, 2015) to explore how sensitive these models are to variations in transition rates.

In some cases, parameter sensitivity analysis reveals that a model’s behavior is controlled by the transition rates of only a few variables, reducing the model’s effective dimension, enabling model reduction and/or reducing the parameter space for more extensive (global) sensitivity analysis and parameter fitting. Based on these results, we suggest that parameter sensitivity analysis should be a part of the construction of a stochastic Boolean model of intermediate size if a detailed mechanistic understanding is important. We provide our MATLAB toolbox *ExaStoLog* as a tool to carry out such analysis for user-defined logical models. The toolbox contains the core calculation method along with various visualization and sensitivity analysis tools and is available on GitHub with a detailed tutorial.

## 2 Materials and Methods

### 2.1 Mathematical derivation

The master equation of a stochastic logical model is a first-order homogeneous system of linear ODEs (Stoll *et al.*, 2012). The state variables of the master equation are the time-dependent probabilities of the 2^*n*^ states of a Boolean model of *n* variables. The solution of such a system could, in theory, be directly calculated by exponentiation of its transition matrix, but since the dimension of the matrix grows exponentially with the number of variables, this is possible only for very small systems.

However, this is not the only possible path of an exact calculation, and the sparsity of the transition matrix (typically less than 0.1% of matrix entries are non-zero) can be exploited to push the limits of an exact calculation in terms of model size. In deterministic chemical kinetics, it was shown (Gunawardena, 2012; Mirzaev and Gunawardena, 2013) that after using timescale separation to eliminate non-linearities, biochemical systems in the mass-action approximation can be described as linear systems of a labeled, directed graph *G* and the linear differential equation

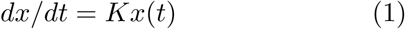

where *x*(*t*) is the concentration of chemical species and *K* the kinetic matrix (called the Laplacian matrix of the directed graph *G* in (Mirzaev and Gunawardena, 2013)).

For such a linear system, as proved in (Mirzaev and Gunawardena, 2013), for any graph *G* and any initial conditions *x*(0), the system always converges to a stable steady state (stationary solution) and this solution can be exactly calculated from the kinetic matrix *K* and the initial condition *x*(0). Mathematically, the master equation of a continuous-time Markov process, of which stochastic Boolean models are examples, is identical to Equation 1, having no non-linearities and guaranteed to have a stable solution, therefore its exact solution can also be obtained. While the dimension of the kinetic matrix *K* grows exponentially with the model’s *n* variables as 2^*n*^, this problem is mitigated by the fact that *K* is very sparse, as shown in Fig. 2A, therefore it does not need to be stored in its full matrix form.

Below we summarize the derivation of the stationary solution.

Linear dynamics is completely determined by the eigenvalues and eigenvectors of the kinetic matrix *K*, and the general solution (Gorban and Radulescu, 2008) to the homogeneous linear equations in 1 is

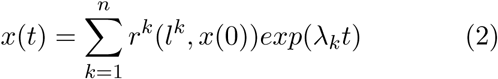

where *λ*_*k*_, *r*^*k*^, *l*_*k*_ are respectively the eigenvalues, the right eigenvectors and the left eigenvectors of the kinetic matrix *K*, with the normalization condition (*l*^*i*^*r*^*j*^) = *δ*_*ij*_, where *δ*_*ij*_ is Kronecker’s delta.

Since here, as usually for Boolean models, only the stationary solution of the model is computed, we do not need to calculate all eigenvectors, but only the kernel (alternatively called the nullspace) of the kinetic matrix. We denote the right kernel matrix *R*, and the left kernel matrix of corresponding left eigenvectors (row vectors) as *L*. *R* contains those right eigenvectors (column vectors) *r*^*k*^ for which *Kr*^*k*^ = 0, which also means that *dx*(*t*)*/dt* = *Kr*^*k*^ = 0. In other words, the right kernel of *K* contains the terminal vertices of the STG, ie. the attractors of the logical model. This means that the right kernel has nonzero elements only in the rows corresponding to states in terminal strongly connected components (SCCs) of the STG. Attractors of a logical model can be individual *stable states* (alternatively called *fixed points*), corresponding to single columns of *R*. In terms of the STG, these states are terminal vertices that have only incoming edges but no outgoing ones: once the model reaches one of these states it cannot escape it. Alternatively, attractors can be *cyclic attractors* comprised of multiple states, appearing as cycles of multiple vertices in the STG with edges within the cycle but no edges leaving it.

We also know that the corresponding eigenvalues for the columns of the right kernel are 0, since there is no further transient dynamics once the system has reached a stable state. Therefore, with the exponential terms being 1, it follows from Equation 2 that the stationary solution of the system, *x*^*^ as *t* → ∞, for any initial condition *x*(0) is:

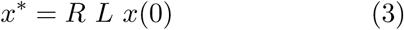

In practice, when obtaining the stationary solution, we do not calculate eigenvectors by standard methods for two reasons. One is the dimension of the kinetic matrix, making these calculations too time consuming. Second, eigenvectors calculated by standard methods from *K* would satisfy the criteria *Kr*^*k*^ = 0 and *l*^*k*^*K* = 0, but not the normalization condition of *δ*_*ij*_ = (*l*_*i*_, *r*_*j*_) and therefore yield numerically incorrect values for *x*^*^.

We can instead build *L* and *R* from the directed graph of the STG, by decomposing its kinetic matrix as follows, described in (Mirzaev and Gunawardena, 2013), a method faster then eigenvector calculations and yielding the numerically correct values for *x*^*^.

If *K* is an *n*×*n* matrix and the dimension of its right nullspace (kernel) has *q* columns, then let us define *R* as a *n* × *q* matrix whose columns are a basis for the column nullspace. Correspondingly, *L* is defined as a *q* × *n* matrix whose rows form a basis for the row nullspace. We do not yet have *L* and *R*, but we know we need to choose them in a way that they satisfy the following conditions:

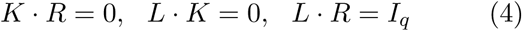

where *I*_*q*_ is an identity matrix of dimension *q* × *q*.

If the conditions in Equation 4 hold then the stationary solution is as in Equation 3, *x*^*^ = *R L x*(0).

The *R* and *L* matrices satisfying the conditions in Equation 4 can be built by decomposing the kinetic matrix *K*, as graphically illustrated in Fig. 1.

**Figure 1:**
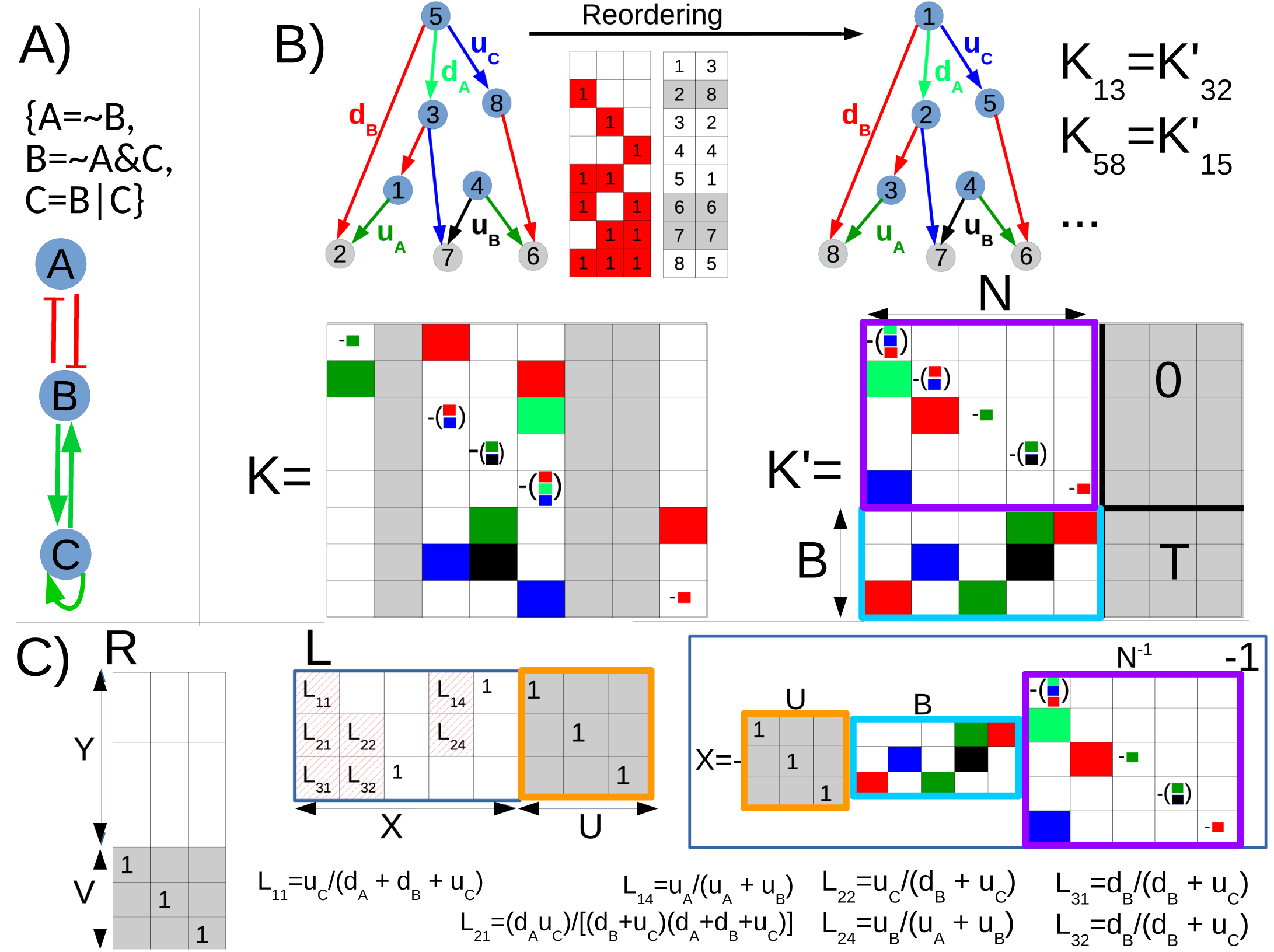
Graphical illustration of the exact calculation method for a 3-node logical model. (A) Influence graph and logical rules of the model (B) Reordering of the state transition graph by topological sorting and the kinetic matrix, moving terminal (attractor) states (2,6,7) in the lower right block of *K*. The colors of the nonzero entries of the kinetic matrix correspond to the transition rates of the model, with the diagonal elements of *K* equal to the sum of the off-diagonal entries in the given column (represented by the smaller squares), taken with negative sign. The red-white heatmap between the two directed graphs (state transition graphs, STG) shows the 8 states of the model with their original and topologically sorted ordering. Terminal vertices corresponding to attractors are in gray. *K* is the original kinetic matrix, *K*′ is the reordered one corresponding to the STG above. *K*′ has a block structure as described in Eq. 5, with terminal vertices moved to the right. (C) Construction of the right and left kernels from the reordered kinetic matrix. R has a number of columns equal to the number of states in terminal SCCs, in this case 3 terminal vertices corresponding to 3 separate stable states. In each column the nonzero element (block *V*) is in the row of the terminal vertices, after reordering these are the last rows 6,7,8. The left kernel *L* is constructed by transposing *V* of the right kernel and then calculating the block *X* from the blocks *B* and *N* of the reordered kinetic matrix as *X* = *−U* · *B* · *N* ^-1^. The nonzero terms of *X, L*_*ij*_ are rational functions in the transition rates of the model, encoding the conservation laws between attractor states and the rest of the state space.

**Figure 2:**
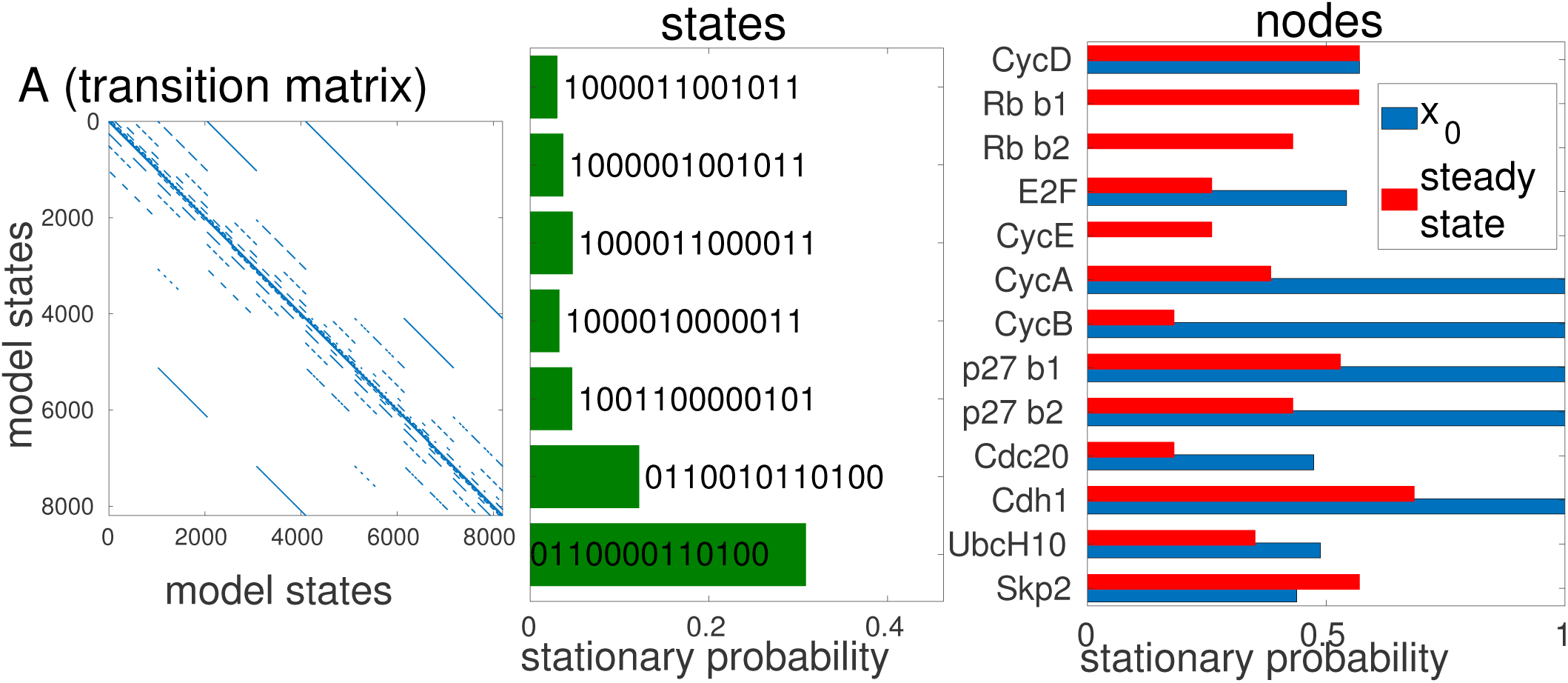
Transition matrix and stationary solution of the mammalian cell cycle model from (Traynard *et al.*, 2016), influence graph shown on SI Fig. 3. (A) Transition matrix *A*. Each axis represents the 2^13^ logical states of the model. Nonzero elements in blue are the transitions between model states. *A* has 2^26^ ≈ 6.7*e*7 entries with only 5.3e4 of them nonzero. The relationship between the transition matrix A and the kinetic matrix *K* is 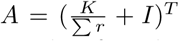, where *r* are the transition rates and *I* is the identity matrix. The element *A*_*ij*_ of the transition matrix is therefore the (normalized) transition rate between states *i* and *j*. (B) Stationary probability values of the model’s attractor states with a probability higher than 3%. The two states at the bottom are the fixed points in one of the two subgraphs of the STG, the other states are part of the cyclic attractor in the other subgraph. (C) Initial and stationary probabilities of model variables.

First, as shown in Fig. 1B for a 3-node logical model, the vertices (states) of the STG need to be topologically sorted and the kinetic matrix *K* reordered accordingly. After topological sorting, the index of a strongly connected component *i* (SCC) is always smaller than that of *j* if there is a directed path from *i* to *j*. The mapping between indices (row and column in the kinetic matrix) and the corresponding logical states (see Fig. 1A) needs to be retained to eventually have the correct assignment of the probability values to model states. The ordering of vertices *within* rings (SCCs of more than one vertex) is of no consequence, therefore the topological sorting is done on the metagraph of the STG, with multi-vertex SCCs treated as single vertices. After sorting the metagraph, the constituent vertices of the multi-vertex SCCs are again unmerged, with their indices having values between the indices of the directly upstream and downstream vertices and their intra-SCC ordering following the initial ordering of the binary states of the logical model. Usually the graph of the STG of logical models has many irreversible transitions, therefore many SCCs are single vertices (as in Fig. 1 where each SCC is a single vertex).

Once this reordering is done (performed in ExaStoLog by the built-in MATLAB function *toposort*), the reordered kinetic matrix (denoted *K*′ in Fig. 1(B)) will have a block structure. The *n - u* columns corresponding to the terminal SCCs are on the right of the kinetic matrix, *u* being the total number of vertices in non-terminal SCCs.

The block structure of K will be the following, as also shown in Fig. 1(B):

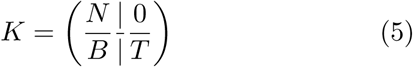

Here and in Eq. 6 horizontal and vertical lines show the borders between blocks and the parentheses the limits of the matrix. In the case of the model in Fig. 1(A), we have 3 terminal vertices, which are all stable states (fixed points), so *T* and the block above it are all zeros, since there are no connecting edges between these vertices or outgoing edges from them.

In the case of cyclic attractors, there are nonzero elements within *T*, representing the edges between the states of the terminal cycle(s), but the block above *T* contains again zeros only.

Now we can construct the right and left kernels from *K*′, which will have the block structure:

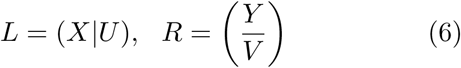

In the right kernel *R*, any row of *R* whose index is not in a terminal SCC contains only zeros, so *Y* = 0 and each column of *V* has nonzero components only for the rows of the terminal SCCs, which are the last rows. In the case of cyclic attractors, a column-wise normalization needs to be applied to respect conservations using the spanning trees of the SCCs as described in (Mirzaev and Gunawardena, 2013), also implemented in the ExaStoLog toolbox. In the case of a model with only stable states as attractors, as in Fig. 1 (C), the nonzero elements of *V* are simply ones.

To build the left kernel, since *L* · *R* = *I*_*n*_, by block multiplication this requires *U* · *V* = *I*_*q*_. To build the block *U*, we transpose *V*, as the left kernel is made up of row vectors, and replace all nonzero elements by 1. In our case, in Fig. 1 (C), these are ones in the right kernel too. *U* · *V* = *I*_*q*_ is therefore satisfied.

The columns of the kinetic matrix sum to zero, 1 · *K* = 0 (with 1 a row vector of ones) and this applies also for its block containing terminal SCCs, and so *U* · *T* = 0. The remaining block of the left kernel, *X* can finally be determined from the fact that *L · K* = 0, which requires by block multiplication that *X* · *N* + *U* · *B* = 0 and since *N* is always invertible after reordering *K*, we obtain *X* = *-U* · *B* · *N* ^-1^.

*N* is a lower triangular matrix if the STG contains no cycles, and contains very few elements in its upper triangular section if there are small non-terminal cycles. Therefore, its inversion is a fast calculation. If the STG of a model contains large (more than a thousand vertices) non-terminal cycles, this inversion can set a limit to our calculation. In the models we have analyzed (Table 1) we have not confronted this problem and we expect most biologically relevant logical models not to have such SCCs in their STG. The calculation of *X* is shown graphically in Fig. 1 (C), with the nonzero terms of *X* in symbolic form.

**Table 1:** Boolean models analyzed in manuscript. Calculation time is for a single solution of one set of initial conditions and parameters, on a CENTOS computer with 8 cores (Intel(R) Xeon(R) CPU X5472 3.00GHz), without parallelization. Besides overall model size, the calculation time depends on the existence of cycles in the STG, as well as the chosen initial conditions, in particular how many of the disconnected subgraphs (due to non-dynamic input nodes) of the STG have states with nonzero initial probability. In parenthesis in the column ‘nodes’ are the numbers of dynamic nodes, excluding input nodes.

| Model | nodes (dynamic) | STG (Mb, density) | calculation time |
| --- | --- | --- | --- |
| Traynard *et al.* (2016) | 13 (12) | 1.4Mb, 7e-4 | 0.35-0.6 sec |
| Zañudo *et al.* (2017) | 20 (16) | 237Mb, 7e-6 | 0.4-3 sec |
| Cohen *et al.* (2015) | 20 (18) | 297Mb, 8e-6 | 2.5-25sec |
| Sahin *et al.* (2009) | 20 (19) | 318Mb, 9e-6 | 2-19 sec |

The nonzero elements of *X* of *L* are rational functions in the model’s transition rates, and it is these terms of the left kernel that encode the conservation laws between the model’s attractors and its nonterminal states. Mathematically they are ratios of polynomials with (only) positive coefficients, originating from the forks and cycles of the STG that distribute the initial probabilities on the vertices of the STG into the attractor states. As shown by Fig. 1 (C), even for a 3-node model, the denominators contain quadratic terms, and for larger models contain polynomials of high order. In other words, the dependence of stationary solutions on transition rates is a complex mathematical expression.

In summary, up to the limit that we can store the transitions of a logical model its stationary solution can always be obtained by topological sorting of its state transition graph and matrix calculations on its reordered kinetic matrix. Using our ExaStoLog toolbox for biological models of around 20 variables the calculation of the stationary solution is of the order of seconds, and the memory requirement is well below 1GB, so the calculations are feasible on a personal computer.

### 2.2 ExaStoLog toolbox: calculation of solutions, visualization and parameter sensitivity analysis

We implemented the above steps of calculating the stationary solution of a logical model in the *ExaStoLog* MATLAB toolbox, available on GitHub. The user first needs to input a logical model in *BoolNet* (Müssel *et al.*, 2010) format using standard logical notation. The generation and topological sorting of the STG and the identification of its cycles are steps independent of the values of the transition rates, therefore these are performed only once for a given model. This is done by the functions *fcn_build_stg_table* and *fcn_scc_subgraphs*. Because the existence of large cycles in the STG is the main limiting factor in the subsequent matrix calculations, *fcn_scc_subgraphs* outputs the size and number of cycles in the STG. The matrix calculations that depend on the numerical values of the transition rates and the initial values are encoded in the function *split_calc_inverse*. The time of calculation and the memory requirement for the STG are shown for four different models that we analyzed in Table 1.

The STG of the (Zañudo *et al.*, 2017) model contains no cycles, therefore its solutions are particularly fast to calculate. The model of (Traynard *et al.*, 2016) has one (terminal) cycle of 270 states, the (Cohen *et al.*, 2015) model a few dozen cycles of 64 to 256 vertices, and the (Sahin *et al.*, 2009) model 32 cycles of 192 vertices and one cycle of 1536 vertices. The calculation times mainly reflect these constraints.

Biological models often have input nodes that are not dynamic, representing environmental conditions such as the presence of a drug or extracellular ligand. Such models have STGs made up of disconnected subgraphs. In this case, the time of calculation depends on whether the states having a nonzero initial probability are in a single or in multiple subgraphs, as shown by the lower and upper bounds in column 4 (calculation time) in Table 1.

Besides the calculation of the stationary solution for an individual set of parameters and initial conditions, ExaStoLog contains 16 other functions to visualize the results and to perform parameter sensitivity analysis and parameter fitting. All figures in the main text except Figure 1 and all figures of the SI except SI Fig. 1, 3-4 and 6-7 were generated by the functions of ExaStoLog. A detailed tutorial showing the use of these functions with examples is available in the GitHub repository.

**Figure 3:**
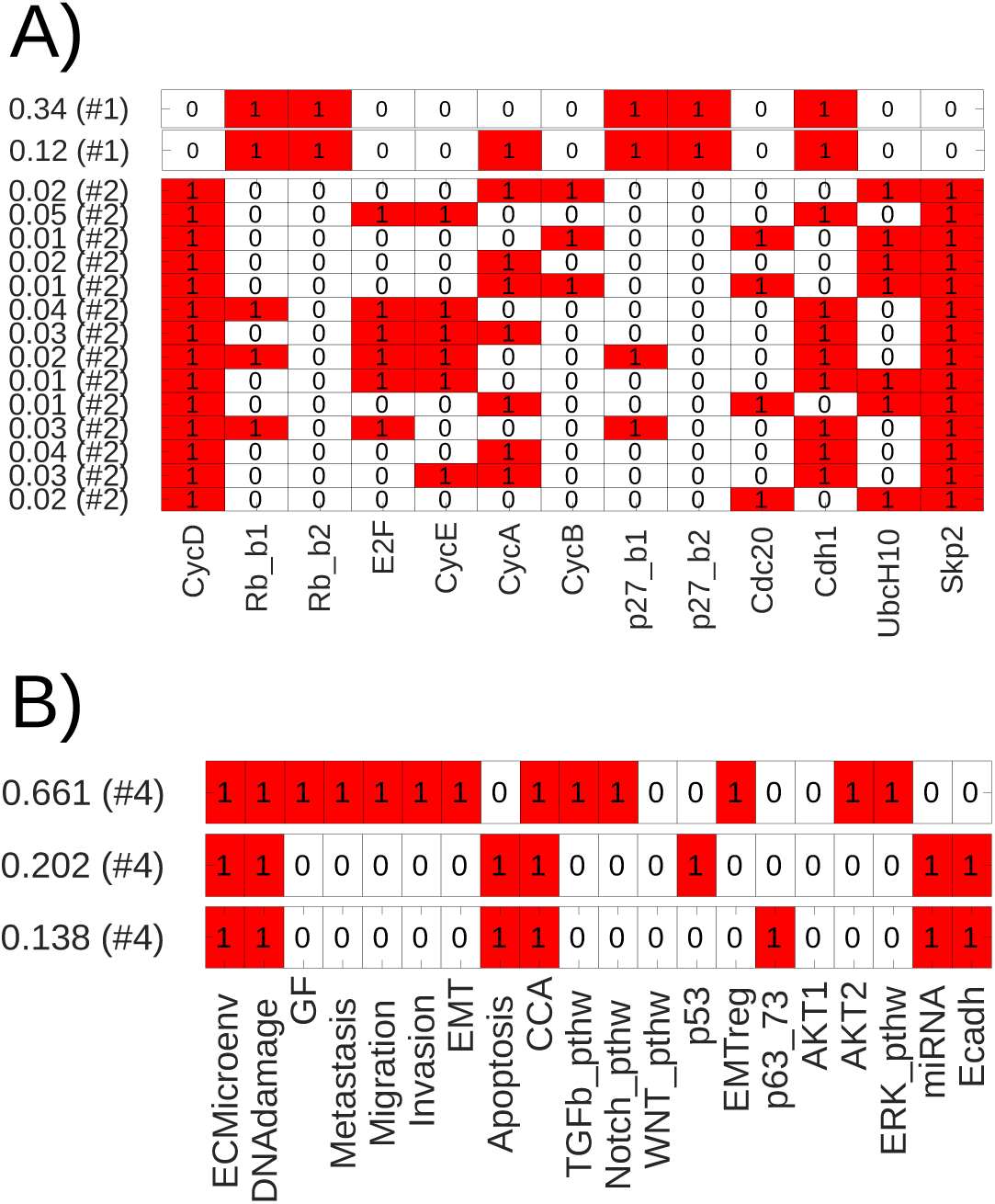
Attractors and their stationary probabilities for two logical models with different types of attractors. Plots by ExaStoLog function *fcn plot statsol bin hmap*. The value to the left of each row is the probability of the given state, with the nodes of the model on the x-axis, red color standing for a variable having the value of 1 (activated). The number in parenthesis is the index of the STG’s subgraph that the attractor state is located in. (A) Attractors of the mammalian cell cycle model from (Traynard *et al.*, 2016). Initial conditions were defined so that the 2 subgraphs of the STG both contain states with nonzero initial probabilities. The two states at the top are separate stable states in subgraph 1 of the STG with CycD=0 (Cyclin D). The lower 13 states, with CycD=1, are part of a cyclic attractor of 270 states (in subgraph 2), of which only those with more than 1% probability are shown. Stable states are automatically plotted with a small gap between them, whereas states in a cyclic attractor are plotted as one block. (B) Attractors of the EMT (epithelial-mesenchymal transition) model from (Cohen *et al.*, 2015). All attractors are separate stable states.

**Figure 4:**
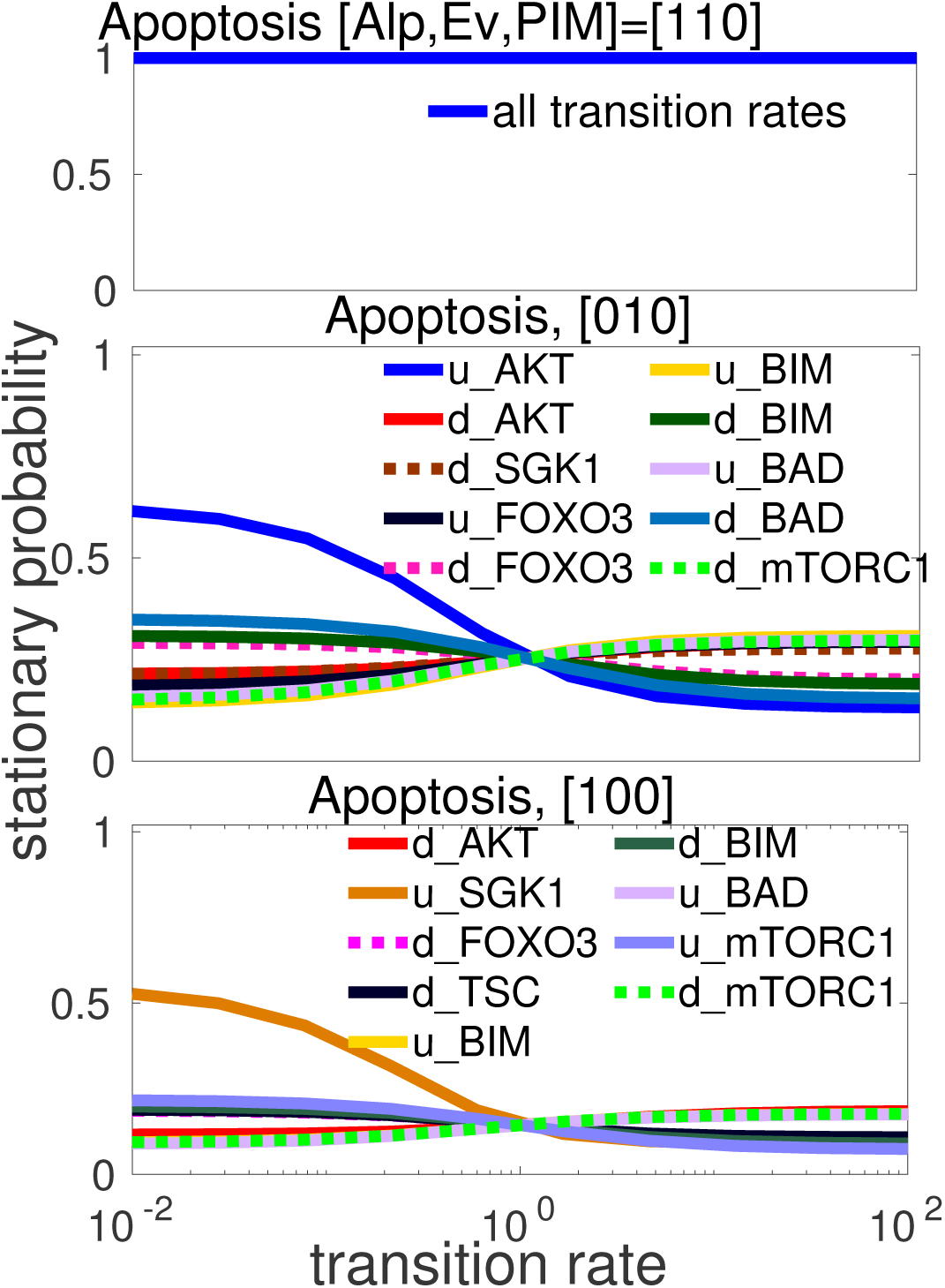
One-dimensional parameter scan of the 20-node Zanudo breast cancer model, with the stationary probability value of the Apoptosis variable (y-axis) shown as a function of the transition rates in the legends. Only the transition rates where the variation in at least one variable’s value is larger than 0.1 are shown. The three panels show the model’s behavior under 3 different initial conditions defined by the value of the input nodes: the drugs Alpelisib and Everolimus and the node PIM (representing members of the PIM protein family). Under double inhibition (and PIM=0), shown on the top panel, the model has a single attractor state with Apoptosis=1, irrespective of the value of transition rates.

To ensure reproducibility and correctness of the results, we first compared the results of multiple models to MaBoSS (Monte Carlo) simulations. We checked models both with separate stable states and cyclic attractors and verified that the results are identical to stochastic simulations, as shown on SI Figure 1 and 2, the latter model of the mammalian cell cycle (Traynard *et al.*, 2016) having a large (270 states) cyclic attractor.

Below we discuss the results obtained by ExaStoLog’s functions for parameter sensitivity analysis for four published Boolean models of different biological processes, listed in Table 1.

## 3 Results

The exact method described above is best-suited to study the stationary solutions and identify the key parameters (transition rates) of logical models of intermediate size, up to 20-25 nodes, depending on memory. We show results with 4 biological models of this size that we selected from the literature to illustrate our exact method and the features of ExaStoLog in terms of sensitivity analysis and parameter fitting. We describe the models in more detail in the SI Section 2, here we give only a very brief summary: (Traynard *et al.*, 2016) presents a discrete model of the mammalian cell cycle. The (Zañudo *et al.*, 2017) model describes the signaling pathways involved in breast cancer and focuses on resistance mechanisms. The (Cohen *et al.*, 2015) model explores the dynamics of the early steps of the metastatic process. Finally, (Sahin *et al.*, 2009) is a Boolean model of breast cancer with an emphasis on ERBB2 overexpression. The model of mammalian cell cycle is known to have a cyclic attractor, testing ExaStoLog’s ability to deal with multi-state attractors, whereas the others are expected to have attractors that are fixed points.

We pose the question to what extent the behavior of these models are parameter-dependent and if parameter sensitivity analysis can aid model reduction, similarly to chemical kinetics (Radulescu *et al.*, 2008).

First, we visualize the model’s attractors (Fig. 2B and Fig. 3), that can be either separate stable states (fixed points) or multi-state cyclic attractors, and their corresponding probabilities. The stationary probabilities of the model variables are simply the probability-weighted sums of these attractor states. These values can be interpreted as the probability that a variable of the logical model is activated. For some models, there are many attractor states. For example, in the case of the breast cancer model (Zañudo *et al.*, 2017), if the model is initiated with uniform initial conditions across all states, so that all the subgraphs of its STG contain states with positive initial probability, there are 39 fixed points, as shown on SI Fig. 5. Others, such as the mammalian cell cycle model (Traynard *et al.*, 2016) have a large cyclic attractor made up of many states. For these cases, looking at the stationary probability value of model *variables* and how these change from their initial value is more biologically informative. The stationary solution by model variables is shown in Fig. 2C for (Traynard *et al.*, 2016) and SI Fig. 5 for (Zañudo *et al.*, 2017).

**Figure 5:**
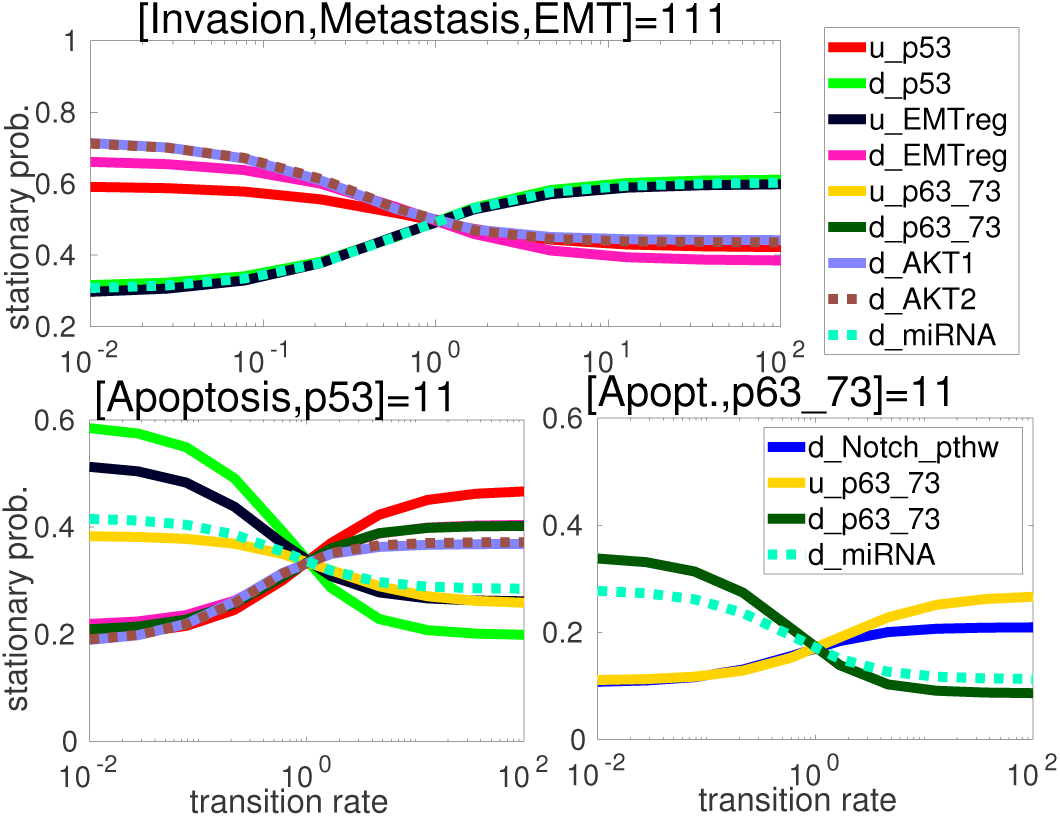
One-dimensional parameter scan of the EMT model from (Cohen *et al.*, 2015), showing the stationary probability (y axis) of the 3 model attractors (see Figure 3B) as a function of the transition rates shown in the legends. The transition rates shown on the top panel and the lower left panel are the same. The input nodes ECMicroenv and DNAdamage were set to 1. The titles above the panels show the variables distinguishing the three states.

The method allows the identification of the transition rates that are more important in defining the model’s behavior than others. The behavior of interest can be for instance the stationary probability of model variables representing biomarkers or phenotypes. We first gauge parameter sensitivity by onedimensional parameter scans. After having scanned all transition rates, those where model variables show significant variation can be selected.

In the case of the (Zañudo *et al.*, 2017) model of drug effects in breast cancer, the sensitivity analysis showed that transition rates have differential effects as a function of the initial value of input nodes. Specifically, as shown in Fig. 4, in the presence of one of the two drugs (and deactivation of PIM), it is the activation (0 → 1) rate of alternatively *AKT* or *SGK1* that has the most potent effect on the stationary probability value of the Apoptosis node. The presence of both drugs, in contrast, invariably leads to 100% Apoptosis, a robust feature of this model.

Alternatively, we can look at the probability of given attractor *states* corresponding to phenotypes, as in Fig. 5 for the (Cohen *et al.*, 2015) model of the early steps of metastasis. For this model, the analysis shows that the decision point between the model’s proliferative-invasive and apoptotic behavior mainly depends on the transition rates of the nodes *p53, p63 73, AKT1, AKT2, Notch pathway* and *miRNA*.

For the cell cycle model, the same analysis shows (SI Fig. 9) that the distribution of probabilities between the two fixed points of this model is determined only by the transition rates *u Rb b2* (*Rb b2* stands for the higher activation level of the retinoblastoma gene) and *d p27 b2* (similarly for the p27 gene). Here, of the large number (270) of states of the cyclic attractor, only 13 show a significant sensitivity to parameters. The others have low probabilities that cannot be significantly amplified by parametric changes. Interestingly, each of these parameter-sensitive states has a single transition rate that can increase their probability up to 40%.

We also found an example of a model’s behavior being robust to relative changes in transition rates: the breast cancer model of (Sahin *et al.*, 2009) has only one attractor state (a fixed point) if all transition rates are nonzero. It is only knockdowns of the model nodes *CDK6, CyclinD1* or *CDK4*, where the initial value and the 1→0 transition rate of the node are set to 0, that make it possible to access the model’s other fixed point, where *pRB* =0, meaning that cell cycle progression (G1/S transition) is blocked. The effects of the different knockdowns for this model is shown on SI Fig. 8.

Besides the stationary values of states or model variables, the local sensitivity (see SI section 3.1.2) to the transition rates can also be visualized. In the case of the EMT model, shown on SI Fig. 10, this analysis reiterates the importance of the transition rates for *p53, p63_73* and the node representing the Notch pathway.

One-dimensional parameter scanning only covers a small subspace of a multidimensional parameter space along its axes. To extend our analysis, based on the results of one-dimensional parameter analysis, the transition rates that have a significant effect on stationary solutions are selected and the multidimensional parameter space of these rates is explored using the Latin Hypercube Sampling (LHS) (Zi, 2011; Constantine and Diaz, 2017) function of ExaStoLog.

The results of LHS are first visualized on scatterplots as shown on SI Fig. 12 with a trendline showing if a variable’s (state’s) stationary probability has a clear trend as a function of particular transition rates. Beyond this visual intuition, the effect of transition rates are statistically analyzed by performing linear regression on the stationary values by the rates (see SI section 3.3.3), calculating and visualizing the coefficient of determination *R*^2^, shown in Fig. 6 (right panel) for the EMT model’s (Cohen *et al.*, 2015) attractor states. For this model, the transition rates of *p53*, and of the nodes representing EMT pathway regulators (*EMT_reg*) and regulatory miRNAs (*miRNA*) have the strongest explanatory value for the stationary solutions.

**Figure 6:**
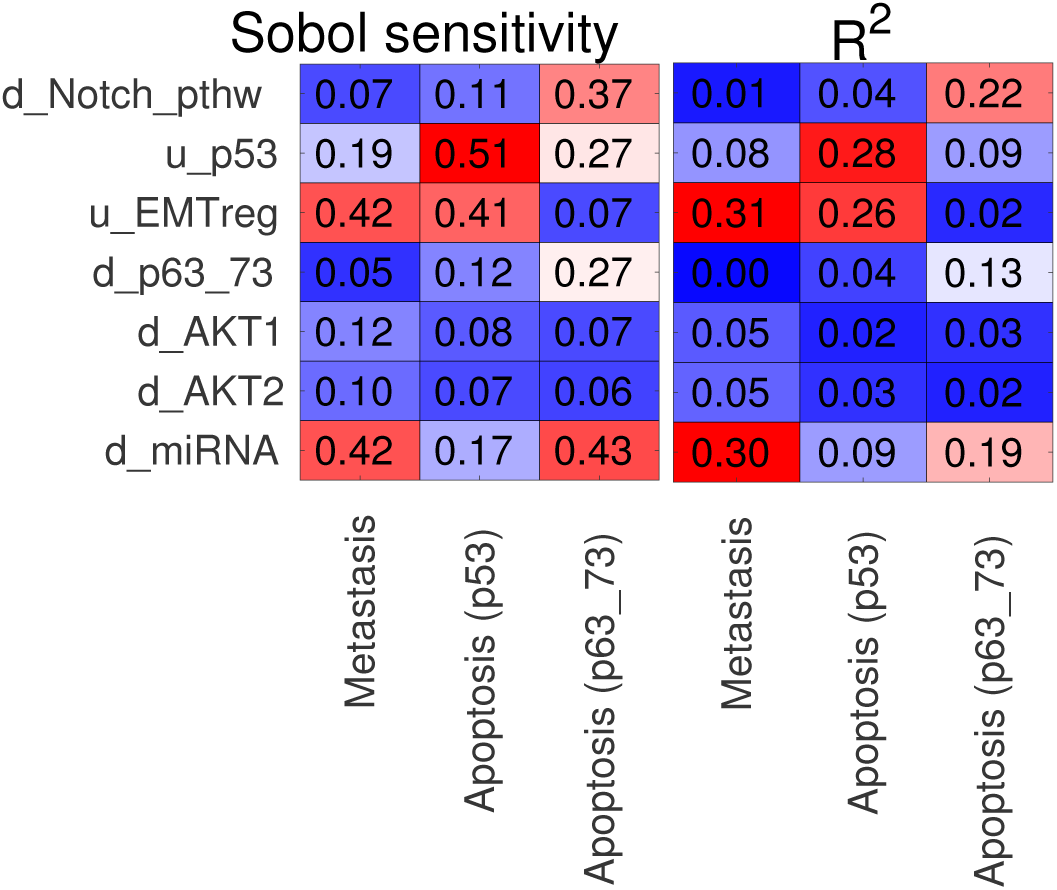
Relative importance of transition rates (y-axis) on the stationary probability values of attractor states (see Fig. 3B) of the EMT model (Cohen *et al.*, 2015), calculated from Latin Hypercube Sampling with 2000 parameter sets. The left panel shows the Sobol sensitivity indices for the transition rates, the sample size was 1000 parameter sets for each resampling. The right panel shows the coefficient of determination (*R*^2^) between transition rates and the probability values of the three states.

Furthermore, calculating correlations between the model’s variables shows (SI Fig. 13) that some variables always have identical values. In this case, these nodes could be merged, reducing the model size. In the case of the EMT model (Cohen *et al.*, 2015), the dominant upstream variables of the *Apoptosis* phenotype seem to be *p53, p63 73, miRNA* and *Ecadh*.

Performing linear regression on the stationary values by the transition rates assumes a monotonic relationship between them, which is not necessarily the case. Sobol sensitivity index (Constantine and Diaz, 2017) is a global sensitivity measure without this limitation that quantifies the relative amount of variance in a variable’s value due to variation of the individual parameters. Calculation of the Sobol sensitivity index, described in SI section 3.3.4, requires recalculating the solutions by individually replacing columns of the matrix of parameters generated by LHS. In the case of the EMT model (Cohen *et al.*, 2015), shown in Fig. 6 (left panel) the Sobol sensitivity indices show a similar pattern as *R*^2^, since there are no nonmonotonic effects.

Finally, if quantitative data is available for a model’s variables or states it is possible with ExaStoLog to perform parameter fitting of the model’s transition rates. If a model’s nodes are proteins, quantitative phosphoprotein data is an ideal data type, often used in another semi-mechanistic modeling approach, modular response analysis (Dorel *et al.*, 2018; Klinger *et al.*, 2013). As described above, the stationary solutions are complex rational functions (ratios of polynomials) of transition rates (Fig. 1C), therefore it is not computationally practical to calculate gradients for parameter fitting for any model larger than a few variables. For this reason, we integrated a gradient-free simulated annealing (Ashyraliyev *et al.*, 2009) algorithm into our toolbox, so that ExaStoLog can be used to directly connect models to experimental data (see SI section 4).

**Figure 7.**
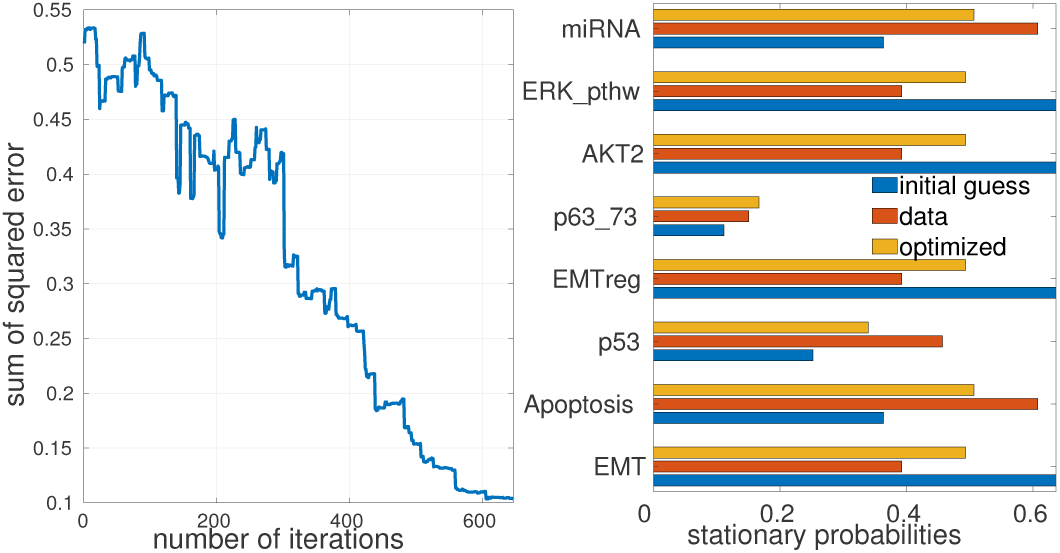
Simulated annealing with 9 transition rates for the Cohen *et al.* (2015) EMT model. The panel on the left shows the convergence process, the panel on the right the data, the initial and optimized values for the model’s variables.

Fitting with simulated annealing of the 20-node EMT model is shown in Fig. 7. Certain models using the initial numerical gradient of the error with respect to the fitted transition rates can be sufficient to perform parameter fitting, with less iterations and therefore lower computation time than for simulated annealing. However neither finding a global minimum nor convergence is guaranteed with this method. The convergence process for the two methods in the case of the EMT model Cohen *et al.* (2015) is shown on Fig. 7 and SI Fig. 16.

With this exact method, it is therefore possible to use stochastic logical models for quantitative modeling and connect models to experimental data directly.

## 4 Discussion

We have shown above that it is possible to calculate the stationary solutions of continuous time stochastic Boolean models by an exact method without resorting to Monte Carlo approximations. Moreover, this exact-continuous calculation method made it easier to explore the question of parameter dependence of this class of models. The examples of Boolean models from the literature have shown that transition rates can indeed have a significant effect on a model’s behavior and typically it is a small subset of all the rates that dominate a model’s behavior, providing a mechanistic understanding of the model.

These findings can be used in several ways. One is the parameterization (instantiation) of models by data, as done in (Béal *et al.*, 2019) to improve the clinically relevant predictive power of Boolean models. In this approach, transition rates can be set to 0 for knockout mutants or their values adjusted based on continuous, such as omics data. Subsequently the phenotypic changes in model behavior can be compared to clinical data to validate the model. *Vice versa*, if data is available on the relative activation level of a model’s variables (or relative frequency of its states (phenotypes)) it can be used to fit transition rates directly.

For example, phosphoprotein measurements with perturbations (Klinger *et al.*, 2013; Dorel *et al.*, 2018; Morris *et al.*, 2011, 2010) can be used to fit transition rates, the same way as it was done with simulated data in Fig. 7. While transition rates are not biochemical constants, they have a plausible biological interpretation as proxies for the timescale of activation-deactivation processes. Similarly, differences in their values across different cell lines or other biological samples can be interpreted as indications for corresponding differences in expression or activity levels of genes, proteins or higher level cellular processes.

## 5 Conclusion

Stochastic logical models represent a powerful framework to study the behavior of cellular networks in terms of their steady state (attractor) behavior and their sensitivity to perturbations. This modeling framework does not require the knowledge of biochemical constants that are often unavailable for most cellular processes. An extension of asynchronously (stochastically) updated, discrete time logical models emerged in recent years (Stoll *et al.*, 2017, 2012) using timescale parameters (transition rates) for the model’s variables to generate continuous time Monte Carlo simulations. This approach produces continuous values (probabilities of activation) for a model’s variables, enabling more quantitative analysis and comparison with continuous biological data.

We took this framework of analysis but implemented it with an exact method adopted from chemical kinetics to perform robustness analysis of several published models. Our analysis confirmed the possibility of efficiently applying exact methods in the context of stochastic logical models, as well as the importance of their parametric analysis, two questions so far neglected in the literature.

This analysis raised several questions that need further investigation. First, for parameter fitting to be more efficient and to ascertain if a global minimum was identified, we need to mathematically analyze the dependence of the stationary solution on transition rates and if its monotonicity can be proved, in which case fitting is a convex problem.

Second, currently our method has a limit in terms of model size due to the memory requirement of explicitly storing the entire state transition graph as a sparse matrix. The first way to mitigate this problem is to store only those states of the STG that are accessible from the states defined by the initial condition and perform the matrix calculations with this reduced transition matrix. This is partially already implemented in ExaStoLog in as much as the sub-graphs of the STG containing no states with positive initial probability are not used for calculations. However the transition matrix is currently not reduced by eliminating inaccessible individual states.

More fundamentally, the indices of nonzero elements of the transition matrix are compositions of arithmetic series defined by the logical rules. This fact could be exploited to avoid simultaneously storing all transitions individually. We are exploring how already existing methods (Bérenguier *et al.*, 2013) to simplify the STG, such as hierarchical transition graphs, can be used in our exact framework. These are again parameter-independent steps therefore they need to be performed only once for a given model, after which individual solutions with different parameter values can be calculated more efficiently.

By making use of some or all of these simplifications we hope to push the limits of our exact method to encompass larger logical models in the future.

## 7 Contributions

M.K. conceived and implemented the method.

M.K. wrote the manuscript with input from A.Z., L.C. and E.B.

V.N. worked on translating the toolbox into Python (ongoing).

## 8 Acknowledgements

We thank our partners Christine Sers, Markus Morkel and Natalie Bublitz at Charité - Universitätsmedizin Berlin for their insightful comments on the biological interpretation of logical models.

