## Supplementary Information for "Exact calculation of stationary solution and parameter sensitivity analysis of stochastic continuous time Boolean models"

Tutorial: https://github.com/mbkoltai/exact-stoch-log-mod/tree/master/doc

#### 1 Comparison of exact method with MaBoSS simulations

The derivation of the stationary states from the full dynamical equations in Section 2.1 of the main text (that we adopted from (Mirzaev and Gunawardena, 2013)) means that the values of the stationary probabilities from stochastic simulations and the exact calculation method must be identical, up to the limit due to noise in Monte Carlo simulations, that increases with a lower number of sample trajectories. To test if our implementation of the exact calculation method is correct, we compared the results from MaBoSS simulations to calculations in our ExaStoLog toolbox (https://github.com/mbkoltai/exact-stoch-log-mod) for several different logical models.

The full dynamics of model state probabilities with a small 3-node model is shown in Figure 1. The MaBoSS input files for this model can be found in the GitHub repository in
model_files/maboss/toymodel_cellfate.bnd,
model_files/maboss/toymodel_cellfate.cfg

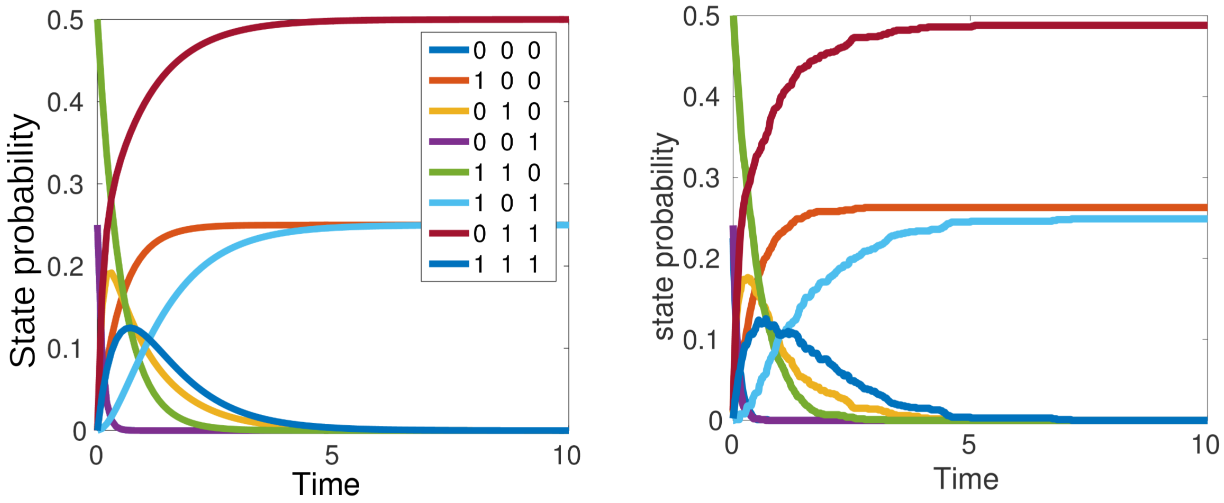


Figure 1: Time course of exact calculation and stochastic simulations of a simple logical model. The dynamics of the probabilities of states of a 3-node logical model with the following rules: $A = A, B = !A, C = B|C$ are shown for the exact calculation (left panel) and stochastic simulations (right). The number of sample trajectories used for stochastic simulations was 1.000. Timecourse calculations are not contained in the released version of ExaStoLog because of their high calculation costs.

We similarly compared results for the cell cycle model of Traynard *et al.* (2016) that has a large cyclic attractor of 270 states, again finding discrepancies not larger than 1%. The MaBoSS files for this simulation are model_files/maboss/traynard2016_mammalian_cellcycle.bnd,
model_files/maboss/traynard2016_mammalian_cellcycle.cfg.

The probability values of states from MaBoSS simulations of a cyclic attractor were averaged over the last 5000 time steps of the simulations to be comparable to the exact solution. We can again see on Figure 2 that the exact method is identical to stochastic simulations up to 1% deviations. We similarly compared ExaStoLog calculations to MaBoSS simulations for other models with fixed point attractors, again getting identical results.

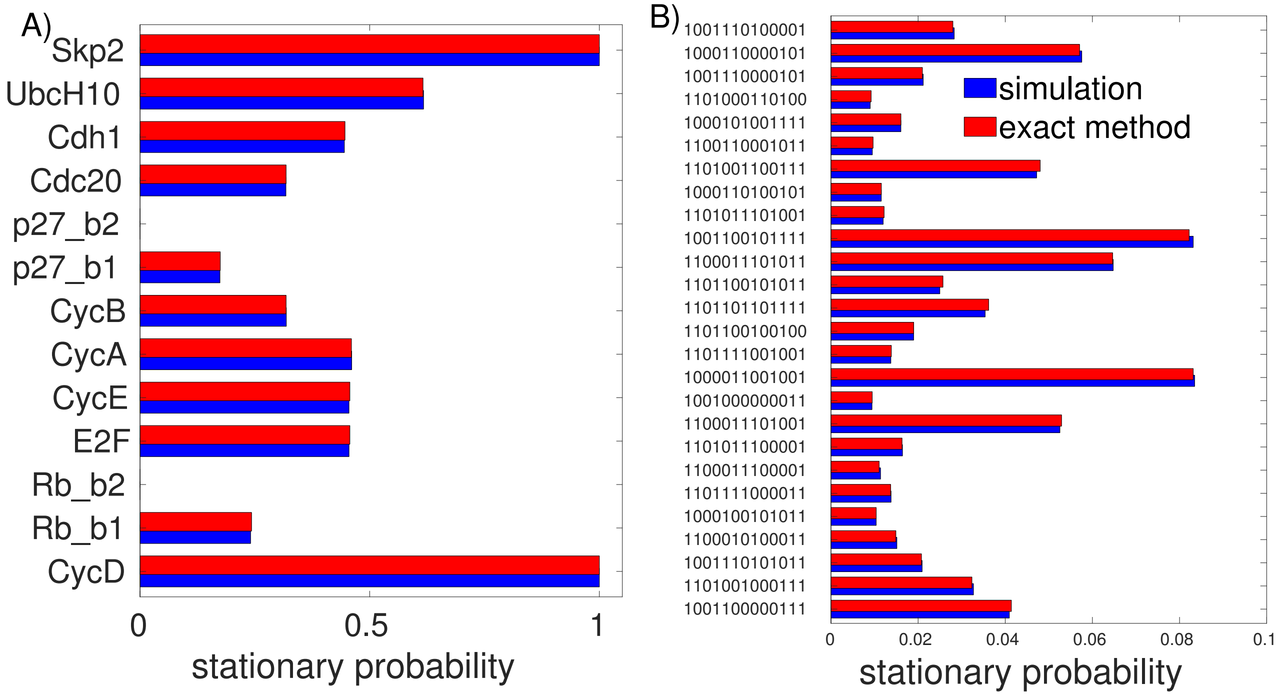


Figure 2: Exact calculation and stochastic simulations of the 13-node mammalian cell cycle model Traynard *et al.* (2016). The stationary probability values for states and (activation of) model variables are shown by the x-axis. The number of sample trajectories for stochastic simulations was 5.000. (A) Probabilities by model variables. (B) Probabilities by states. Only states with at least 1% probability are shown.

### 2 List of models analyzed

#### 2.1 Traynard 2015 mammalian cell cycle model (13 nodes)

Source file at GitHub repository: model_files/mammalian_cc.bnet

This is a 13-node model of the mammalian cell cycle, published in Traynard *et al.* (2016), its influence graph shown on SI Fig. 3. The model has two separate subgraphs. In one subgraph there are 2 fixed point attractors (where Rb_b2, p27_b2=1 and CycD, Skp2=0, the value of CycA differentiating the two states), in the other a large cyclic attractor of 270 states. We analyzed if the stationary probabilities of the two fixed point attractors and the states within the cyclic attractor are sensitive to the transition rates. Results of parameter scans are shown on SI Fig. 9.

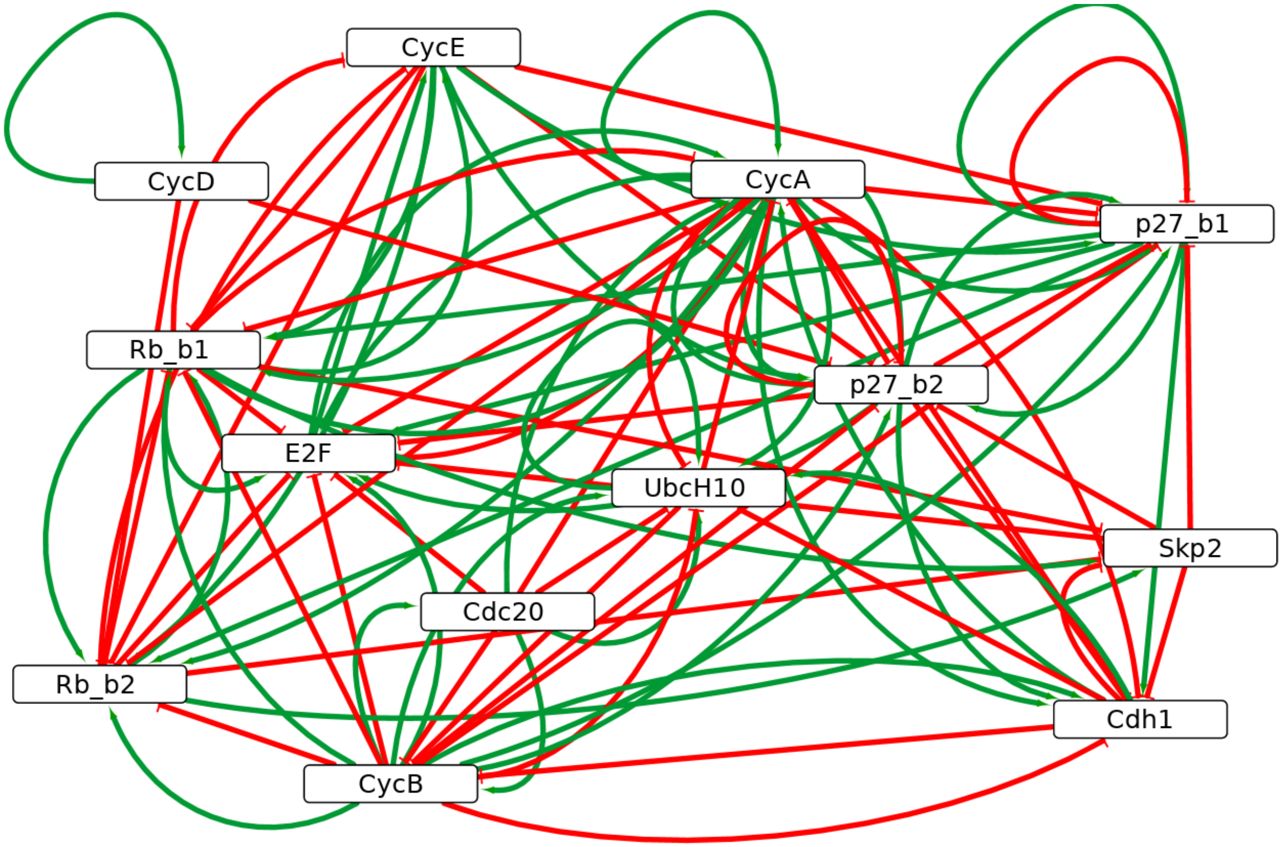


Figure 3: Influence graph of 13-node model of mammalian cell cycle, from Traynard *et al.* (2016)

#### 2.2 Zanudo 2017 breast cancer model (20 nodes)

Source file at GitHub repository: model_files/breast_cancer_zanudo2017.bnet

This model is the 20-node subnetwork of the full model published in Zañudo *et al.* (2017), shown on Figure 5A of Zañudo *et al.* (2017) in the main text. The influence graph is shown on SI Fig. 4. The model has 4 input nodes: Alpelisib, Everolimus, PDK1, PIM, and the state transition graph is made up of 36 disconnected subgraphs. Due to the high number of disconnected subgraphs if we populate all of them by uniform initial conditions we have a large number of attractor states, shown on SI Fig. 5, all of which are separate stable states. The calculation time (see Table 1 in main text) depends on the choice of initial conditions and how many of the subgraphs are populated. Calculation for one subgraph is below one second. The model has two phenotypic nodes: *Proliferation* and *Apoptosis*. We analyze the dependence of the stationary probabilities of these two variables under different initial conditions in terms of the value of Alpelisib, Everolimus and the PIM node, as shown on Figure 4 of the main text. The initial values of these input nodes can be interpreted as the presence of the two drugs and the mutational status of the PIM family of oncoproteins.

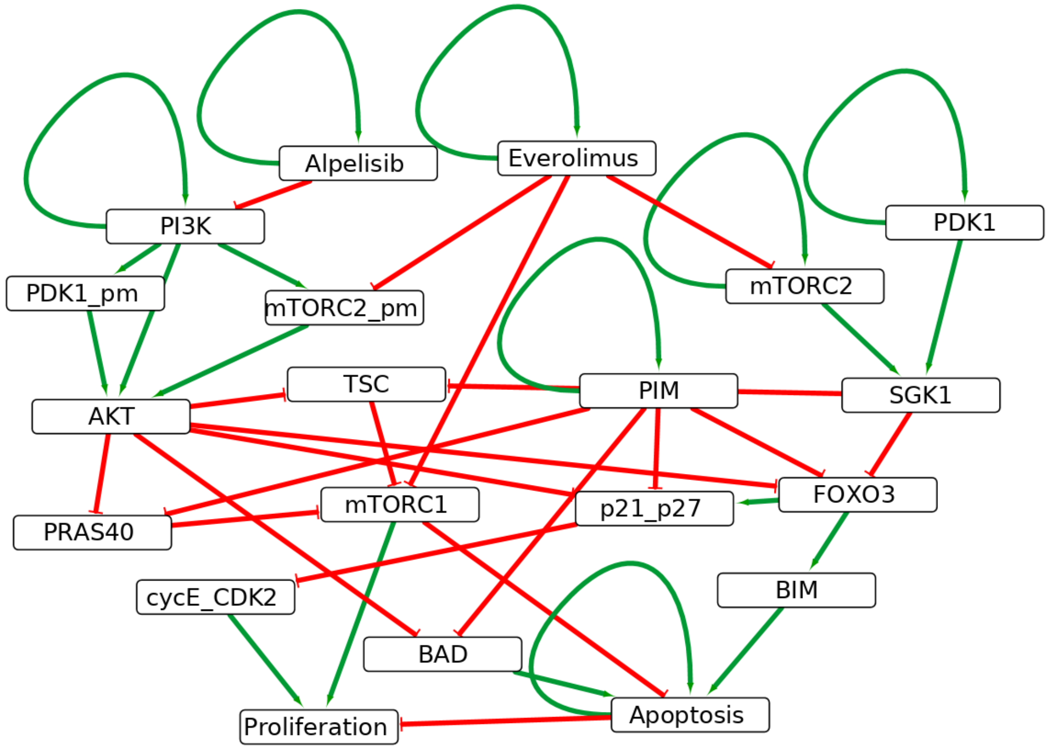


Figure 4: Influence graph of 20-node model of drug resistance in breast cancer, from Zañudo *et al.* (2017)

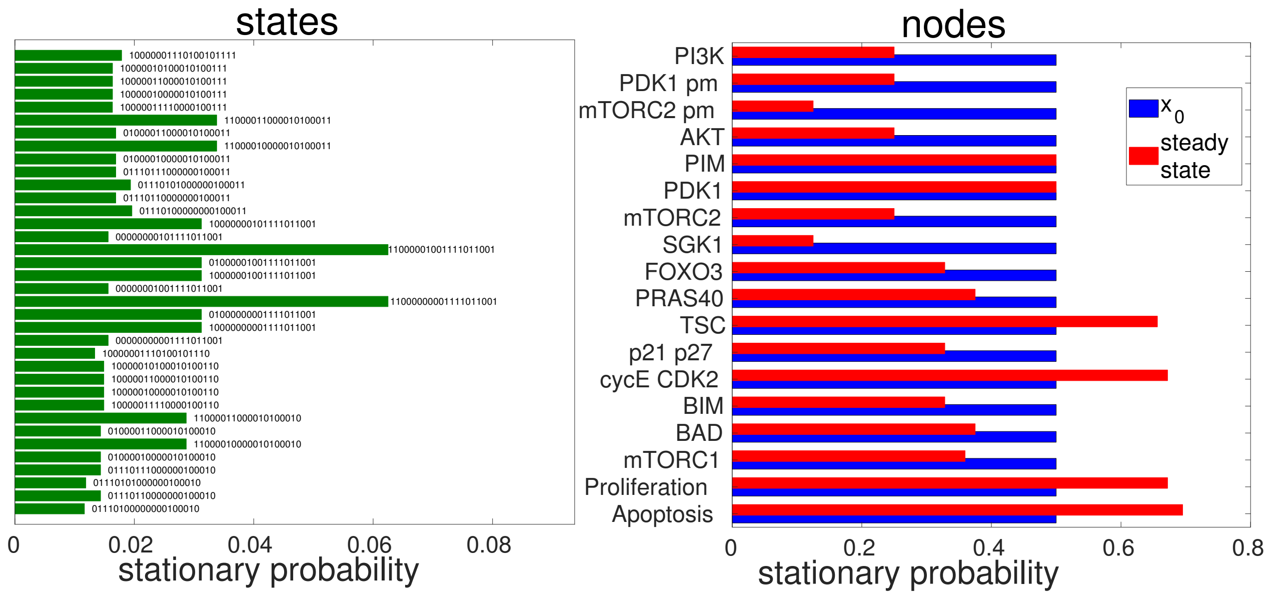


Figure 5: Stationary probability values of states and variables of the breast cancer model from Zañudo *et al.* (2017) with uniform initial conditions across all model states, so all subgraphs of the STG have their states with a positive uniform initial probability.

#### 2.3 Cohen 2015 EMT model (20 nodes)

Source file at GitHub repository: model_files/EMT_cohen_ModNet.bnet
This is the 20-node modularized version of the full model of epithelial-mesenchymal transition (EMT) from

Cohen *et al.* (2015). The influence graph is shown on SI Fig. 6. The model has two groups of phenotypic nodes:
- *Apoptosis* and *CCA* (cell cycle arrest)
- *EMT*, *Invasion*, *Migration* and *Metastasis*

There are two input nodes: *DNAdamage* and *ECMicroenv*. We analyzed the parameter-dependence of the attractor states, characterized by co-activation of one group of phenotypic nodes under the 4 different initial conditions. The results of the sensitivity analysis are summarized on Figures 5, 6 of the main text as well as SI Fig. 10. We have learned from the sensitivity analysis that it is primarily the transition rates d_miRNA, u_p53, d_p63_73, u_EMTreg and d_Notch_pthw that determine the cell fate decision between apoptosis and EMT/proliferation/invasion. Also, we have recapitulated a known (see Cohen *et al.* (2015)) synergistic effect of p53 loss-of-function and Notch pathway gain-of-function mutations in a continuous sense by our two-dimensional scan shown on SI Fig. 11.

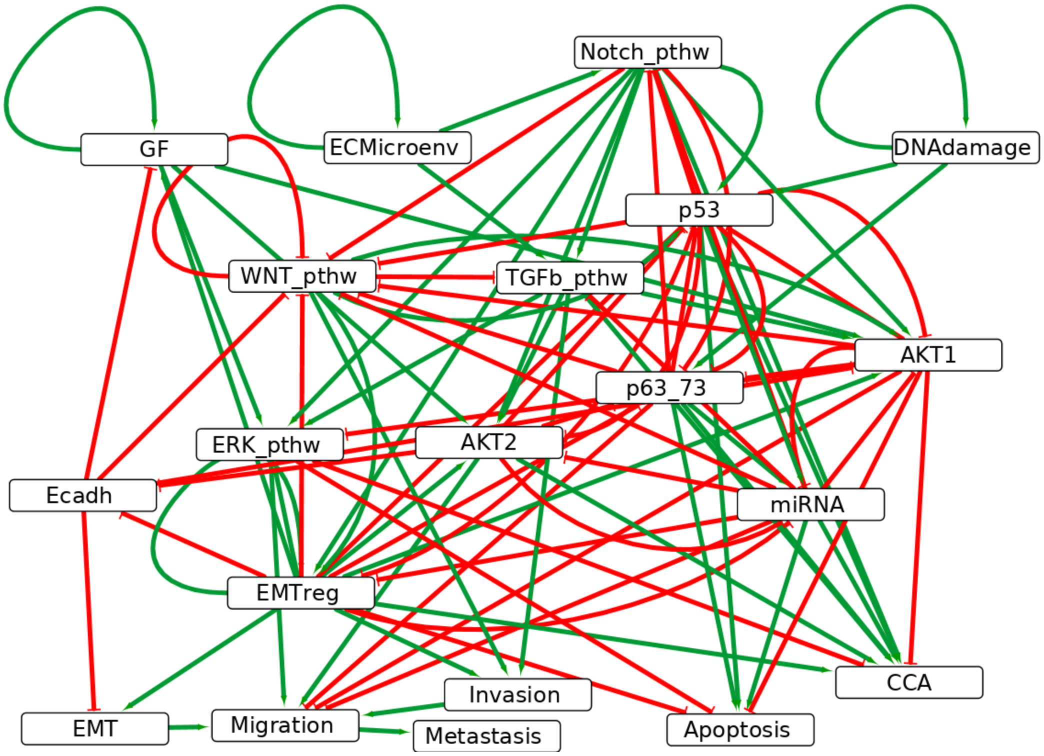


Figure 6: Influence graph of the 20-node model of EMT and invasion from Cohen *et al.* (2015)

#### 2.4 Sahin 2009 breast cancer ERBB-model (20 nodes)

Source file at GitHub repository: model_files/sahin_breast_cancer_refined.bnet
This is a 20-node model on the role of ERBB receptor overexpression/mutations and its inhibition by trastuzumab (a monoclonal antibody) in breast cancer from Sahin *et al.* (2009). The influence graph is shown on SI Fig. 7. The model's output node is *pRB* that represents the phosphorylation of the retinoblastoma protein, that is a proxy for G1/S transition (cell cycle progression) and proliferation. This model's attractor is robust to changes in the values of transition rates: it has only one attractor if all transition rates have a positive value. We simulated knockdowns by setting the initial value and the 'upward' (0-¿1) transition rate *u_node* of nodes to 0. For some knockdowns there is another attractor with *pRB=0*, meaning that the G1/S transition (and hence proliferation) cannot occur. These knockdowns are shown on SI Fig. 8.

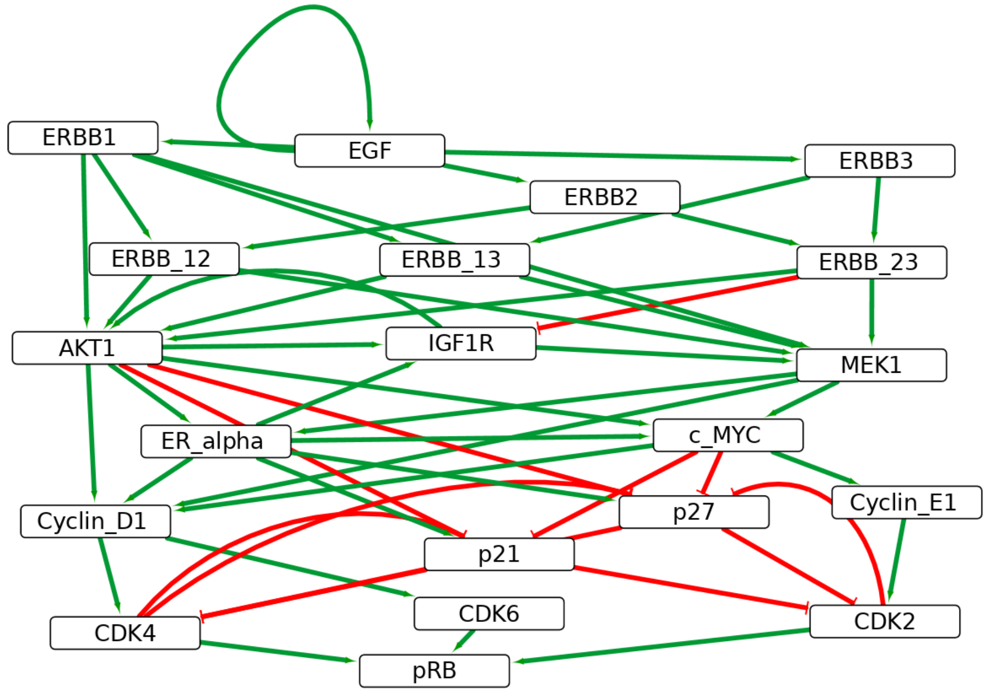


Figure 7: Influence graph of the 20-node model of the role of ERBB receptor in breast cancer from Sahin *et al.* (2009)

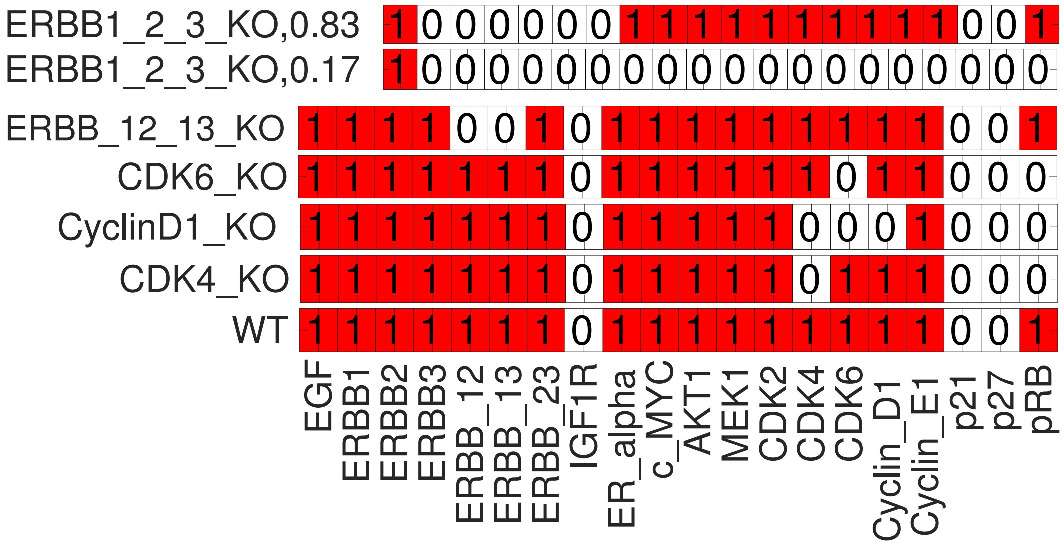


Figure 8: Effects of knockdowns in the 20-node model of breast cancer from Sahin *et al.* (2009)

### 3 Parameter sensitivity analysis

#### 3.1 One-dimensional parameter scans

##### 3.1.1 Values of states/variables as a function of transition rates

We can generate lineplots of parameter scans grouped by the transition rates with the function *fcn_onedim_parscan_plot_by_para* as shown below for the mammalian cell cycle model. This plot can be produced either for the value of model states (SI Fig. 9) or variables. On each subplot we have the stationary probability of those states or variables that show variation above a user-defined threshold as a function of the given transition rate.

With the function *fcn_onedim_parscan_plot_parsensit* we plot the one-dimensional parameter scan with the values of one state/variable on each subplot as a function of the transition rates, for example on Fig. 4 and 5 of the main text.

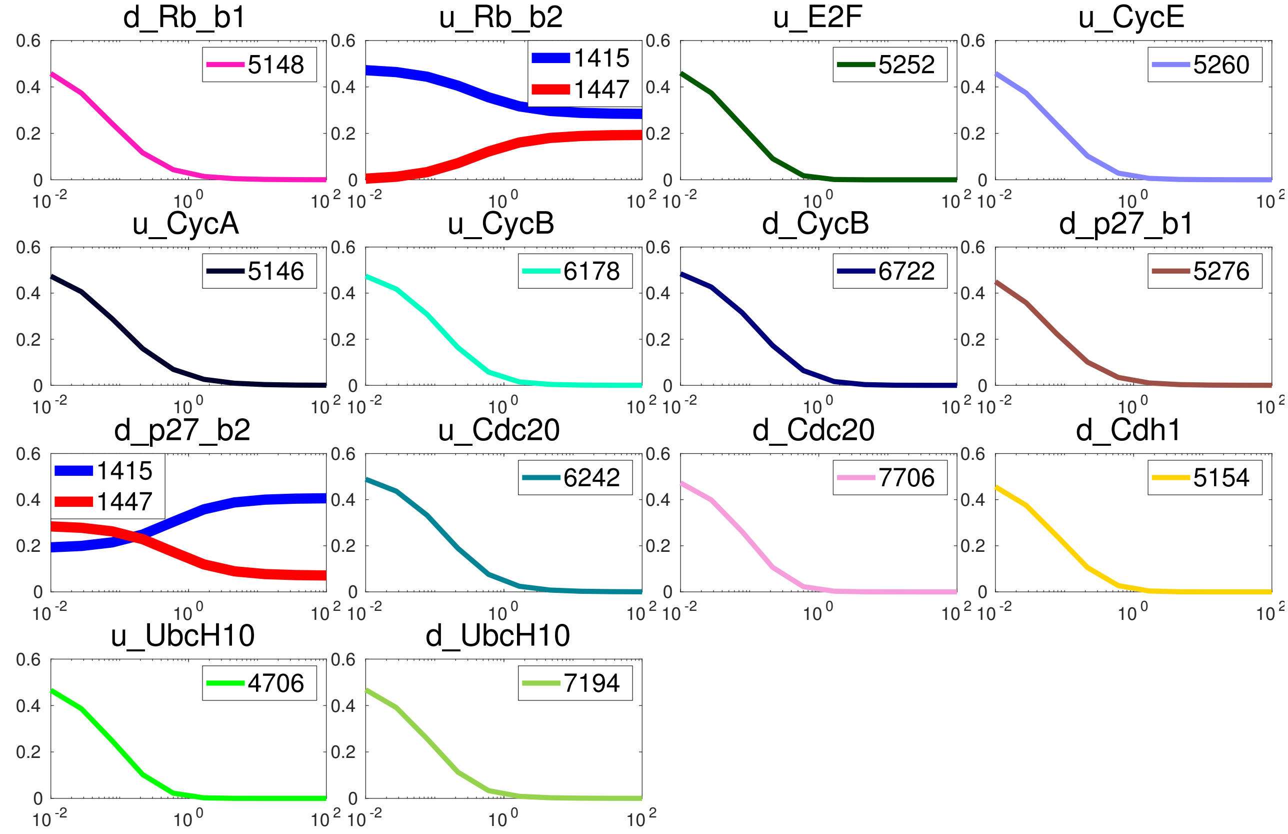


Figure 9: One-dimensional parameter scan for the states of the mammalian cell cycle model, Traynard *et al.* (2016). Each subplot is for one transition rate (values on x-axis), with states showing at least 15% change in their probability (y-axis) shown. The states that are separate fixed points (1415,1447) are shown with thicker lines (panel 2 and 9). All other states are part of the cyclic attractor of 270 states. Legends show the index of the states, with a numbering going from [0000000000000] to [1111111111111].

###### 3.1.2 Local sensitivity of variables to transition rates

Besides the stationary value of the model's states/nodes it can be informative to plot the local sensitivity of the stationary solutions to the transition rates by visualizing their response coefficients, a metric from metabolic control analysis Kacser *et al.* (1995). The response coefficient $R^x_p$ of a variable $x$ with respect to a parameter $p$ is its derivative normalized by the local value of the variable and the parameter:

$$R^x_p = \frac{\partial log x}{\partial log p} = \frac{\partial x/x}{\partial p/p} = \frac{\partial x}{\partial p}\frac{p}{x} \tag{1}$$

As we do not compute stationary solutions symbolically we do not have the symbolic derivatives, but they can be numerically approximated by $\frac{\partial x}{\partial p}\frac{p}{x} \approx \frac{\Delta x}{\Delta p}\frac{p}{x}$ This metric tells us how a variable reacts in relative terms to a small change in a parameter. If $R^x_p = 1$ (-1) this means a 1% change in $p$ induces a 1% increase (decrease) in $x$.

On SI Fig. 10 we show the response coefficients for the EMT model, highlighting stronger responses for the parameters *d_p63_73*, *u_p53* and *d_Notch_pthw* consistently with Fig. 5 of the main text that shows the values of the same variables.

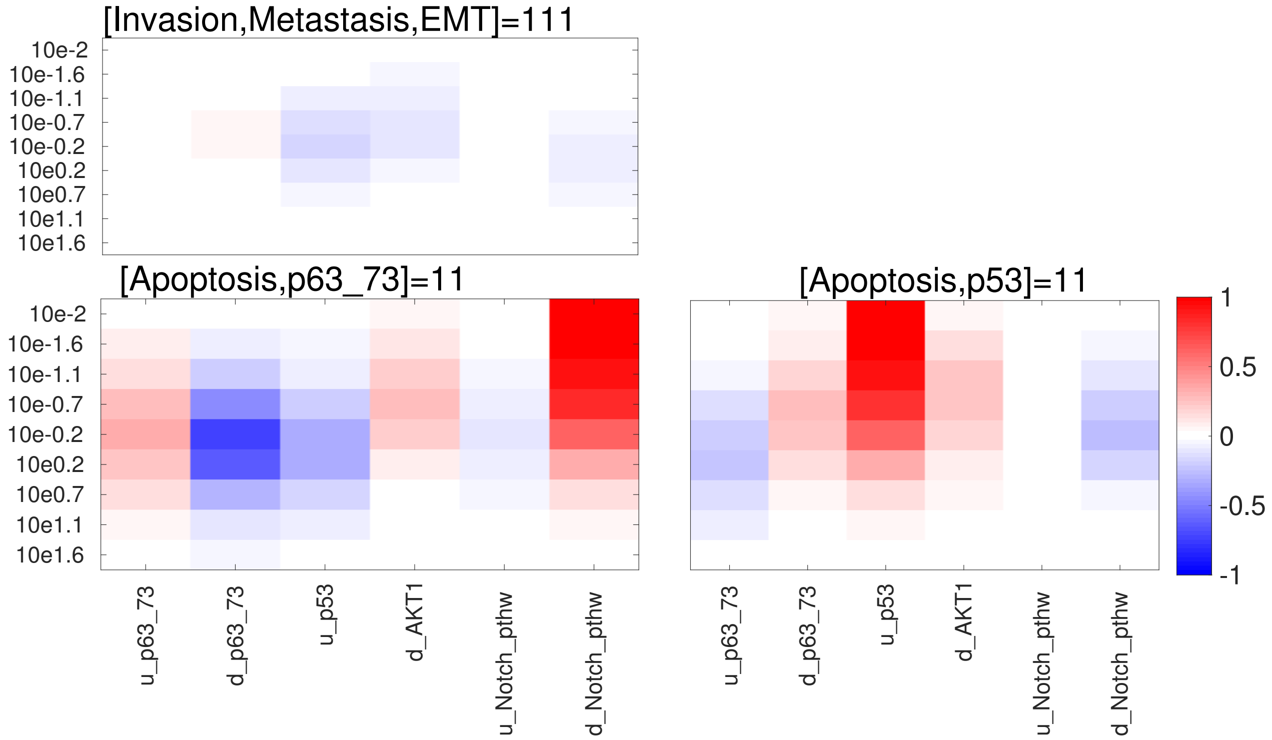


Figure 10: Response coefficients from one-dimensional parameter scan of the attractor states (title above subplots) of the EMT model. The transition rates are shown on the x-axis of the bottom row of subplots, the y-axis shows the values covered by the parameter scan. The color code, going from -1 to 1 shows the value of the response coefficient of the given variable with respect to the transition rate.

##### 3.2 Multidimensional parameter sensitivity analysis with uniform resolution

With the function *fcn_calc_paramsample_table* we can perform multidimensional parameter sampling with a uniform resolution. We recommend to do this in two dimensions, for easy visualization and since otherwise it can be computationally expensive, eg. if the number of value for each parameter is 5, for n parameters we need $5^n$ calculations. For two dimensions we can visualize the results by the function *fcn_plot_twodim_parscan*. Synergistic effect between transition rates can be visualized, as in SI Fig. 11 for *u_p53* and *u_Notch_pthw* with respect to Metastasis in the EMT model of Cohen *et al.* (2015).

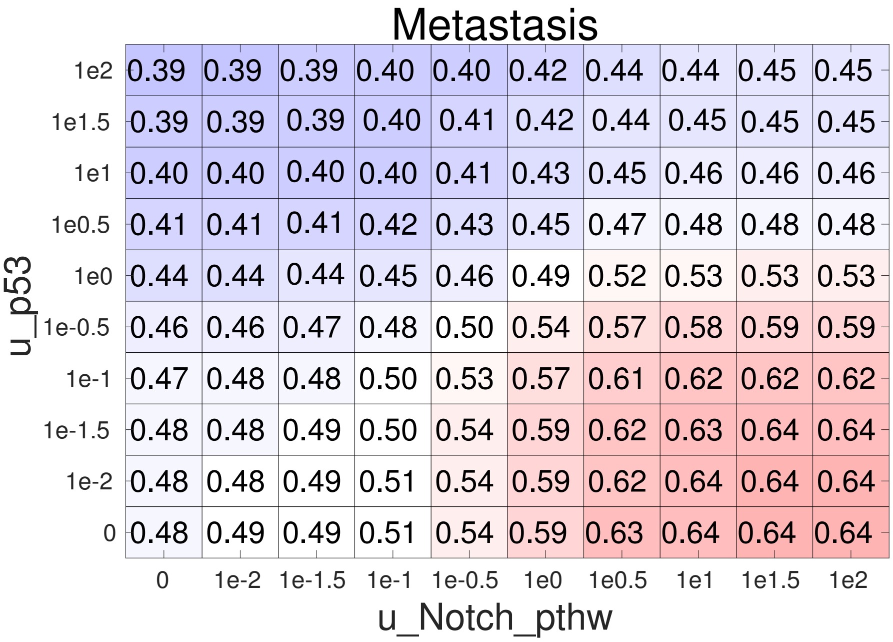


Figure 11: Two-dimensional parameter scan of the EMT model Cohen *et al.* (2015), with the stationary probability value of the Metastasis model variable shown as a function of the transition rates *u_p53* and *u_Notch_pthw*.

##### 3.3 Multidimensional parameter sensitivity analysis by Latin Hypercube Sampling (LHS)

Latin Hypercube Sampling (LHS) is a method to efficiently explore a high-dimensional parameter space. In ExaStoLog we can perform LHS by the function *fcn_multidim_parscan_latinhypcube*, defining the sample size, as well as the type, mean and standard deviation of the distribution we sample from. From one-dimensional parameter scans (with the function *fcn_onedim_parscan_plot_parsensit*) we have the transition rates where there is significant variation in the stationary solutions and perform LHS for only these. Results of the LHS are analyzed visually and statistically by the functions described below.

###### 3.3.1 Scatter plots of LHS

With the function *fcn_multidim_parscan_scatterplot* we can visualize the stationary probability values (of model states or variables) from LHS as a function of transition rates as a scatterplot. The mean value of the stationary probabilities is shown by the red trend line. To calculate the mean the user needs to select the number of bins across the range of values for transition rates.

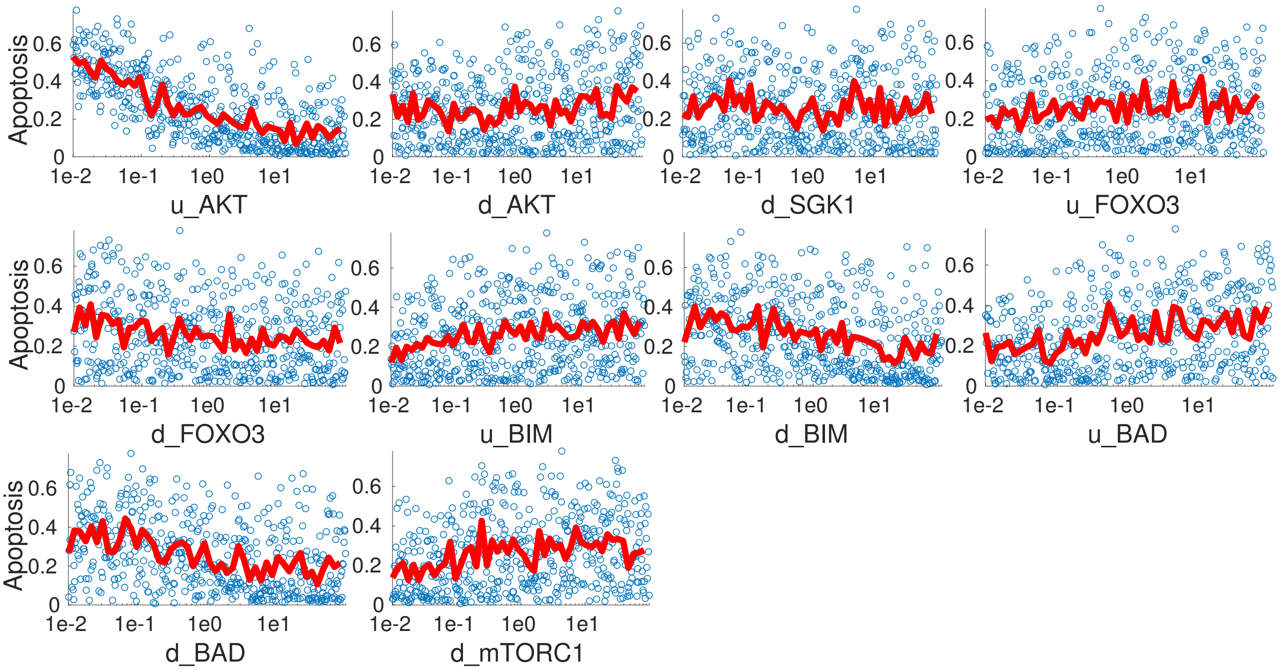


Figure 12: Result of Latin Hypercube Sampling for selected transition rates for the Zañudo *et al.* (2017) breast cancer model. The stationary value of the *Apoptosis* node is shown, with the red line showing its mean value in 50 bins. The visually discernible trends for *d_BAD* and *u_AKT* are confirmed by Sobol sensitivity analysis shown on SI Fig. 15

###### 3.3.2 Correlations between stationary probability values of model variables

Pearson correlation coefficients between the stationary probability values of variables following LHS for the Cohen *et al.* (2015) model are shown on SI Fig. 13. The calculation and plotting is by the function *fcn_multidim_parscan_parvarcorrs* in ExaStoLog. This metric is useful for model reduction: some variables show 100% correlation that suggests that these can be merged into single variables.

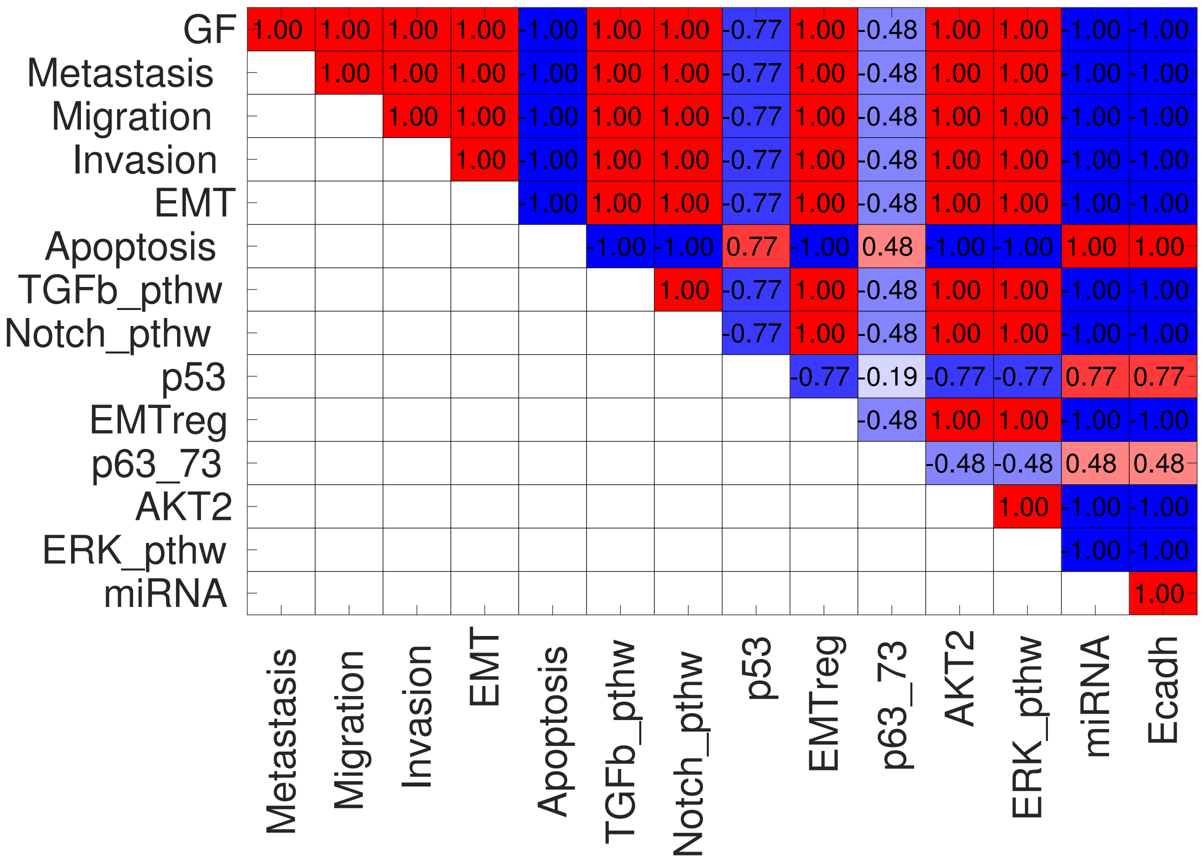


Figure 13: Correlation coefficients between stationary probability values of model variables from LHS for the model Cohen *et al.* (2015)

##### 3.3.3 Coefficient of determination ($R^2$) between the transition rates and the stationary probability values of nodes/states

With the function *fcn_multidim_parscan_parvarcorrs* we can also perform linear regression on the stationary probability values of model states/variables as a function of the transition rates. In the case of lognormal sampling for the LHS, it is recommended to do the regression as a function of the logarithm of transition rates (a built-in feature of the function). The $R^2$ value for each parameter-variable/state pair indicates to what extent the given transition rate predicts the stationary value. Values for the Cohen *et al.* (2015) model are shown on SI Fig. 14. This measure assumes a monotonic effect of transition rates on the stationary probability values, which is usually, but not necessarily the case. An alternative measure that does not have this problem is the Sobol sensitivity index. In the case of transition rates with a monotonic effect these two measures show similar, but not identical values as shown on Fig. 6 in the main text.

##### 3.3.4 Sobol sensitivity index

Sobol sensitivity index is a global parameter sensitivity index that also quantifies nonlinear and non-monotonic effects of parameters on variables. Following LHS the Sobol sensitivity index is calculated by splitting the matrix of randomly generated transition rates from the LHS into two and in one of the sub-matrices replacing one column (corresponding to a transition rate) with newly generated random values. Following that we recalculate the stationary solution of the model. The Sobol sensitivity index quantifies how much of the total variance in a variable is due to variation in the transition rate that was changed. The formulas for the usual numerical approximation are described in Constantine and Diaz (2017). A comparison of the Sobol sensitivity index with $R^2$ values can be seen on Fig. 6. The result of Sobol sensitivity analysis

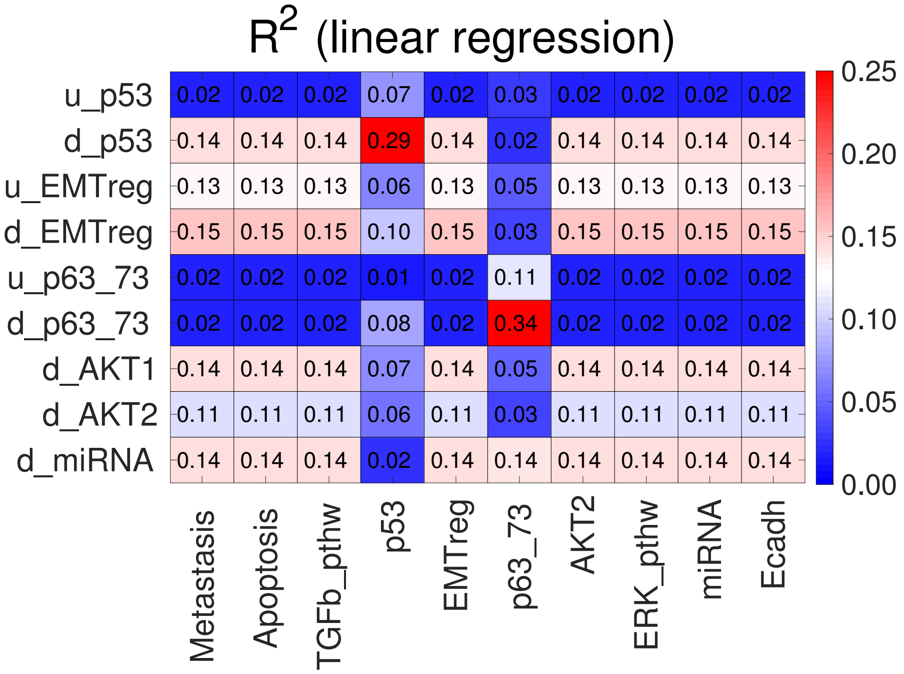


Figure 14: $R^2$ values between model variables and transition rates for the model Cohen *et al.* (2015). Variables that have identical values with those on the plot are not shown. Also transition rates with no or negligible effects were omitted.

for the Zañudo *et al.* (2017) breast cancer model's Apoptosis variable is shown on SI Fig. 15, with $u\_AKT$ and $d\_BAD$ dominating apoptotic behavior.

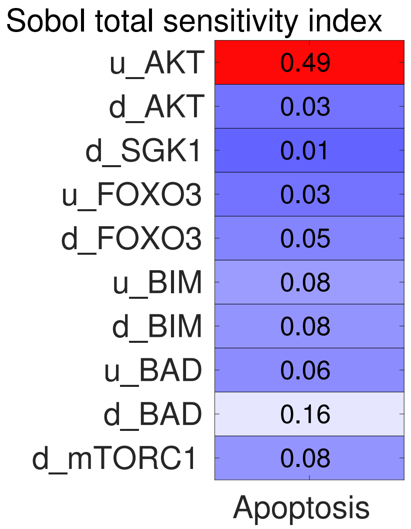


Figure 15: Sobol sensitivity index of transition rates of the Zañudo *et al.* (2017) breast cancer model with respect to the model's Apoptosis variable.

### 4 Parameter fitting

We can fit the stationary probability value of either the model's attractor states or its variables, the latter being linear combinations of the former. With the function *fcn_handles_fitting* of ExaStoLog we create anonymous functions (*fcn_statsol_sum_sq_dev*, *fcn_statsol_values*) to calculate the sum of squared error (SSE) of the value of the model's states/variables with respect to a vector of values (data) provided by the user. The transition rates that we want to fit are selected by the user.

#### 4.1 Parameter fitting by simulated annealing

We use a simulated annealing algorithm *anneal* from MATLAB Central, with some modifications of the script for ExaStoLog (see Tutorial on GitHub). Simulated annealing is a gradient-free, probabilistic parameter fitting method. It often shows slow convergence, requiring thousands of iterations (recalculations of the stationary solution). An example is shown on Fig. 7 of the main text. Results from parameter fitting are plotted by the function *fcn_plot_paramfitting*.

#### 4.2 Parameter fitting by numerical gradient descent

For the models we analyzed in this paper the effect of transition rates on the stationary probability value of model variables is monotonic. Therefore in some cases we can perform parameter fitting by calculating an initial gradient for the sum of squared error as a function of transition rates and reduce the SSE down to a defined level by incrementing the transition rates by this initial direction. At least in the case of some models, such as the Cohen *et al.* (2015) EMT model, this method is faster and leads to a larger error reduction than simulated annealing, as shown by SI Fig. compared to Fig. 7 of the main text. This method is implemented in the function *fcn_num_grad_descent*, that automatically stops (after 2 steps) if the error starts to grow or stagnate. Plotting is with the same function as for simulated annealing, *fcn_plot_paramfitting*.

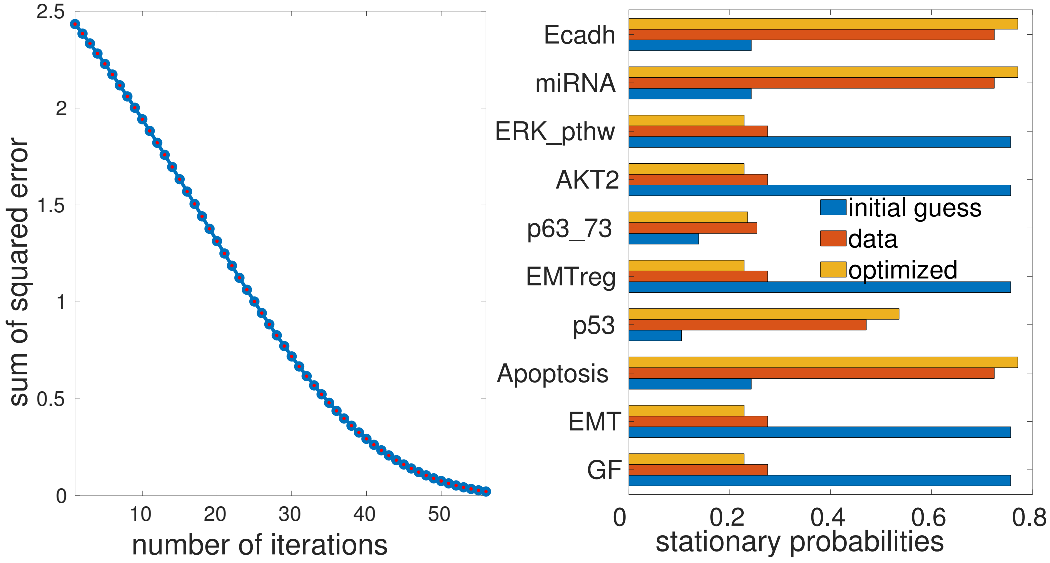


Figure 16: Parameter fitting by numerical gradient descent for the Cohen *et al.* (2015) EMT model. The panel on the left shows the convergence process, the panel on the right shows the data, the initial and the fitted values of the model's variables.
